# Optimization of NLS Composition Improves CRISPR-Cas12a Editing Rates in Human Primary Cells

**DOI:** 10.1101/2022.02.01.478672

**Authors:** Kevin Luk, Pengpeng Liu, Jing Zeng, Yetao Wang, Stacy A. Maitland, Feston Idrizi, Karthikeyan Ponnienselvan, Lihua Julie Zhu, Jeremy Luban, Daniel E. Bauer, Scot A. Wolfe

## Abstract

Type V CRISPR–Cas12a systems are an attractive alternative nuclease platform for specific genome editing applications. However, previous studies demonstrate that there is a gap in overall activity between Cas12a and Cas9 in primary cells. Here we describe optimization to the nuclear localization signal composition and architecture of Cas12a to facilitate highly efficient targeted mutagenesis in mammalian cell lines (HEK293T, Jurkat, and K562 cells) and primary cells (NK cells and CD34+ HSPCs), regardless of Cas12a ortholog. A 3xNLS Cas12a architecture resulted in the most robust editing platform. The improved editing activity of Cas12a in both NK cells and CD34+ HSPCs resulted in pronounced phenotypic changes associated with target gene editing. Lastly, we demonstrated that optimization of the NLS composition and architecture of Cas12a did not decrease the specificity of editing in HEK293T and CD34+ HSPCs. Our new Cas12a NLS variant provides an improved nuclease platform for therapeutic genome editing.

## Introduction

Clustered regularly interspaced short palindromic repeats (CRISPR)–Cas12a is a type V CRISPR-Cas system that has been well characterized and harnessed by the research community for genome editing.^1–3^ The Cas12a system has several unique characteristics that distinguish Cas12a from the more commonly utilized Cas9 nuclease platform from *Streptococcus pyogenes* (SpyCas9). First, the most commonly employed Cas12a nucleases from *Acidaminococcus sp* (Asp) and *Lachnospiraceae bacterium* (Lba) recognize a T-rich (TTTV [V = A/G/C]) protospacer adjacent motif (PAM) sequence and utilize a single ~42 nucleotide CRISPR RNA (crRNA) for target site recognition.^1,3,4^ Additionally, Cas12a cuts distally from its PAM sequence in a staggered fashion, generating a double strand break (DSB) with four or five nucleotide 5’-overhangs, whereas Cas9 cuts proximal to its PAM, typically generating a blunt ended DSB.^1,3,5^ Furthermore, due to Cas12a nucleases’ ability to process CRISPR arrays, multiplex genome editing is more accessible.^6–8^ These properties, along with the favorable precision of Cas12a nuclease platforms, make it an attractive alternative nuclease platform for genome editing applications to the Cas9 system for certain therapeutic and research applications.^9–11^

AspCas12a and LbaCas12a are the most widely employed Cas12a nucleases in the genome editing field due to their promising editing activity in a number of biological systems including fruit flies, mammalian cells, mouse embryos, zebrafish, and plants.^1,4,9,10,12–23^ However, our own studies have shown that wild-type Cas12a nucleases may display lower editing rates than SpyCas9 in primary cells.^14^ Efforts have been made to increase the overall editing activity of Cas12a.^13,24–29^ Notably, enAspCas12a is a recently described engineered Cas12a nuclease with increased editing activity relative to AspCas12a and LbaCas12a at canonical-TTTV PAMs.^25^ Additionally, the enAspCas12a variant is able to efficiently utilize a broader range of PAM sequences (TTYN, VTTV, TRTV, and others) [N = A/C/T/G, Y = C/T, R = A/G].^25^ However, enAspCas12a is more promiscuous, editing a higher fraction of near cognate sequences than wild-type AspCas12a.^25^ Efforts have been made to address the specificity issues of enAspCas12a by introducing mutations to improve the fidelity of DNA cleavage.^25^ Although prior work has made significant progress to improve the efficiency of Cas12a nuclease platforms, these methods are typically focused on engineering specific Cas12a orthologs, as opposed to general approaches applicable to all variants.^24,25,27,30,31^ In this work, we sought to define modifications to Cas12a that are broadly applicable and that would produce improved genome-editing activity for research and therapeutic genome-editing applications without sacrificing its high intrinsic specificity.

Previously, we sought to improve SpyCas9 gene editing in quiescent primary cells by optimizing the sequence composition and the number of nuclear localization signal (NLS) sequences.^32^ We found that SpyCas9 bearing three NLSs (1 N-terminal and 2 C-terminal) [3xNLS-SpyCas9] substantially improved editing activity in primary hematopoietic stem cells.^32,33^ Additionally, we found that Cas12a bearing two C-terminal NLSs (2xNLS-SV40-NLP Cas12a) (NLP = Nucleoplasmin NLS) (Fig. 1A) had improved activity in primary hematopoietic stem cells, transformed cell lines, and zebrafish.^1334^ While this modification to Cas12a increased nuclease activity compared to previous variants of the nuclease, Cas12a proteins with two NLS sequences did not achieve the same level of targeted mutagenesis as 3xNLS-SpyCas9 in CD34+ hematopoietic stem and progenitor cells (HSPCs) at therapeutically relevant targets.^14^ These observations prompted further investigations to improve Cas12a editing activity in primary cells with a focus on examining the impact of the number and composition of NLS sequences on Cas12a ribonucleoproteins (RNP) indel frequencies. To improve the nuclear import of our existing 2xNLS-SV40-NLP Cas12a nuclease, we substituted the SV40 Large T-antigen NLS, herein referred as SV40 NLS, with a more efficient import sequence, c-Myc NLS, and added a third NLS to either N- or C-terminus of Cas12a (Fig. 1A). Here, we show that the modifications to the NLS framework of Cas12a improved editing activity, achieving editing efficiencies that approach 100% target sequence disruption. The three NLS C-terminal variant of Cas12a resulted in markedly increased knockout efficiency without compromising the high specificity of Cas12a in transformed cell lines (HEK293T, Jurkat, and K562 cells), NK cells, and CD34+ HSPCs. Taken together, these new modifications to the NLS framework of Cas12a provide improved editing frameworks for both studying gene function and therapeutic genome editing in primary cells.

**FIG 1.**
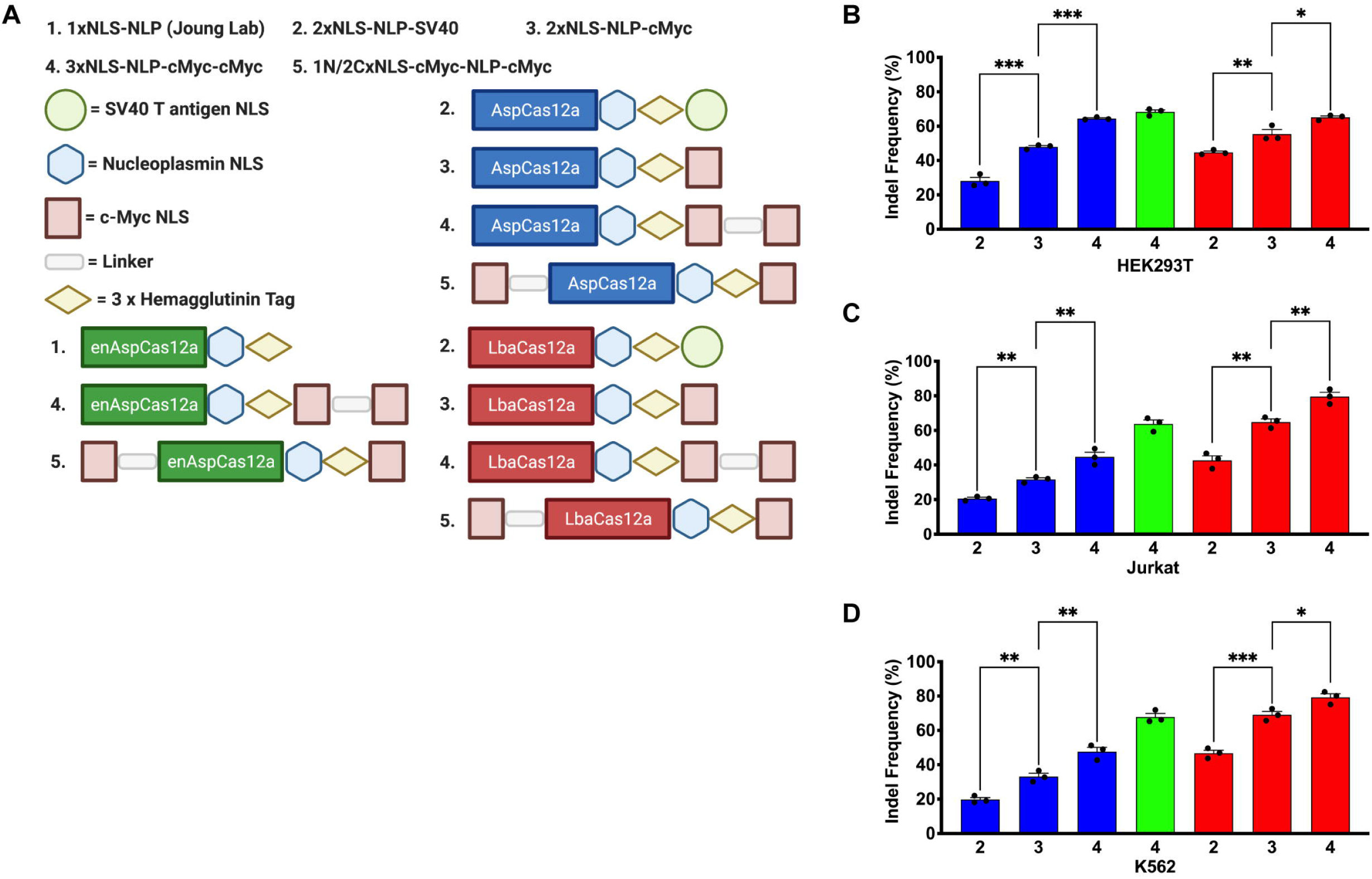
New Cas12a NLS variants demonstrate improved activity in transformed cell lines. (A) Schematic representation of the prior designs for Asp/LbaCas12a (2xNLS-NLP-SV40) and enAspCas12a (1xNLS-NLP) in addition to the new Asp/enAsp/LbaCas12a NLS framework variants described in this study.^13,14,25,34^ (B) Quantification of editing efficiency by Cas12a RNP targeting *AAVS1* site 1 in HEK293T cells at 5 pmol of protein:crRNA complex when delivered by nucleofection. (C) Quantification of editing efficiency by Cas12a RNP targeting *AAVS1* site 1 in Jurkat cells at 5 pmol of protein:crRNA complex when delivered by nucleofection. (D) Quantification of editing efficiency by Cas12a RNP targeting *AAVS1* site 1 in K562 cells at 5 pmol of protein:crRNA complex when delivered by nucleofection. Results were obtained from three independent experiments and presented as mean◻±◻SEM. *\*P*◻<◻0.05, *\*\*P*◻<◻0.01, \**\*\*P*◻<◻0.001 by one-way ANOVA with Tukey’s multiple comparisons test between each NLS variant. Cas12a ortholog is denoted by color from 1a. Each NLS architecture is denoted by a number from 1A.

## MATERIAL AND METHODS

### Plasmid Constructs

Cas12a nuclease experiments for Neon transfection in cell culture employed the following plasmids: All AspCas12a, enAspCas12a, and LbaCas12a protein expression for protein purification utilized pET-21a protein expression plasmids (Novagen) (Figure 1).^1,25^ AspCas12a, enAspCas12a, and LbaCas12a NLS variant expression constructs were constructed containing a 6xHis tag at the C-terminus for affinity purification. Sequences of NLSs are in Supplementary Table S1. Amino acid sequences for new Cas12a NLS variants are in Supplementary Table S2. Plasmid expression vectors for the key constructs are available from Addgene.

### Transformed Cell Culture and Nuclease Assays

Human Embryonic Kidney (HEK293T) cells were cultured in high glucose DMEM with 10% FBS and 1% Penicillin/Streptomycin (Gibco) in a 37°C incubator with 5% CO_2_. K562 and Jurkat cells were cultured in RPMI 1640 medium with 10% FBS and 1% Penicillin/Streptomycin (Gibco) in a 37°C incubator with 5% CO_2_. These cells were authenticated by University of Arizona Genetics Core and tested for mycoplasma contamination at regular intervals. For Neon transfection, we used early to mid-passage cells (passage number 5-15). Cas12a RNPs were delivered to HEK293T, Jurkat or K562 cells by nucleofection. AspCas12a, enAspCas12a, or LbaCas12a proteins were complexed with the desired crRNA at a molar ratio of 1:2.5 in Neon R buffer (Thermo Fisher Scientific) and incubated at RT for 15 minutes. For HEK293T cells, the Cas12a RNP complex was then mixed with 1 × 10^5^ cells in Neon R buffer at the desired dosage and electroporated using Neon® Transfection System 10 L Kit (Thermo Fisher Scientific) using the suggested electroporation parameters: Pulse voltage (1150v), Pulse width (20ms), Pulse number (2). For Jurkat and K562 cells, the Cas12a RNP complex was then mixed with 1 × 10^5^ cells in Neon R buffer at the desired dosage and electroporated using Neon® Transfection System 10 L Kit (Thermo Fisher Scientific) using the suggested electroporation parameters: Pulse voltage (1600v), Pulse width (10ms), Pulse number (3).

### AspCas12a and LbaCas12a Protein purification

Protein purification for all AspCas12a, enAspCas12a, or LbaCas12a NLS variants used a common protocol as previously described.^13^ The plasmid expressing each Cas12a protein was introduced into *E. coli* Rosetta (DE3) pLysS cells (Sigma) for protein overexpression. Cells were grown at 37°C to an OD600 of ~0.2, then shifted to 18°C and induced for 16 hours with IPTG (1 mM final concentration). Following induction, cells were pelleted by centrifugation and then resuspended with Nickel-NTA buffer (20 mM TRIS + 1 M NaCl + 20 mM imidazole + 1 mM TCEP, pH 7.5) supplemented with HALT Protease Inhibitor Cocktail, EDTA-Free (100X) [ThermoFisher] and lysed with M-110s Microfluidizer (Microfluidics) following the manufacturer’s instructions. The protein was purified with Ni-NTA resin and eluted with elution buffer (20 mM TRIS, 500 mM NaCl, 250 mM Imidazole, 10% glycerol, pH 7.5). Cas12a protein was dialyzed overnight at 4°C in 20 mM HEPES, 500 mM NaCl, 1 mM EDTA, 10% glycerol, pH 7.5. Subsequently, Cas12a protein was step dialyzed from 500 mM NaCl to 200 mM NaCl (Final dialysis buffer: 20 mM HEPES, 200 mM NaCl, 1 mM EDTA, 10% glycerol, pH 7.5). Next, the protein was purified by cation exchange chromatography (Column = 5ml HiTrap-S, Buffer A = 20 mM HEPES pH 7.5 + 1 mM TCEP, Buffer B = 20 mM HEPES pH 7.5 + 1 M NaCl + 1 mM TCEP, Flow rate = 5 ml/min, CV = column volume = 5ml). For Cas12a variants chosen for in-depth analysis of editing activity, cation exchange chromatography was followed by size-exclusion chromatography (SEC) on Superdex-200 (16/60) column (Isocratic size-exclusion running buffer = 20 mM HEPES pH 7.5, 300 mM NaCl, 1 mM TCEP). The primary protein peak from the SEC was concentrated in an Ultra-15 Centrifugal Filter Ultracel −30K (Amicon) to a concentration of between 50 to 100 mM. The purified protein quality was assessed by SDS-PAGE/Coomassie staining to be >95% pure.

### Synthesis of human genome specific CRISPR RNAs

Synthetic AspCas12a and LbaCas12a CRISPR RNAs (crRNAs) targeting *AAVS1* site 1, *EMX1* site 1, IFNG, CD96, EnAspTS2 and *DNMT1* site 3 were made by Integrated DNA Technologies (IDT) with their proprietary modifications at each end of the crRNA (AITR1 on 5’ end and AITR2 on 3’ end) (Supplementary Table S3). Synthetic SpyCas9 single guide RNAs (sgRNAs) targeting SpyTS1 and Spy1617 were made by IDT with their proprietary modifications at each end of the sgRNA (Supplementary Table S3).

### Target site indel frequency analysis in mammalian cells by deep sequencing

Library construction for Illumina deep sequencing is modified from our previous report.^13,35^ For analysis of mammalian cell culture experiments, cells were harvested to extract genomic DNA with GenElute Mammalian Genomic DNA Miniprep Kit (Sigma). Briefly, regions flanking each target site were PCR amplified using locus-specific primers bearing tails complementary to the TruSeq Illumina adapters as described previously.^13^ 25-50ng input genomic DNA is PCR amplified with Q5 High Fidelity DNA Polymerase (New England Biolabs): (98°C, 15s; 65°C 30s; 72°C 30s) x 30 cycles. 1 μl of each PCR reaction was amplified with barcoded primers to reconstitute the TruSeq adaptors using the Phusion High Fidelity DNA Polymerase (New England Biolabs): (98°C, 15s; 64°C, 25s; 72°C, 25s) x 10 cycles. Approximately equal amounts of the products (determined by band intensity on gel) were pooled and gel purified. The purified library was deep sequenced using a paired-end 150bp Illumina MiniSeq cassette. MiniSeq data analysis for indel frequencies at on-target and off-target sites was performed using CRISPResso2 software (*CRISPRessoBatch --batch_settings --amplicon_seq -- ignore_substitutions --guide_seq --cleavage_offset 1 --plot_window_size 25 -- window_around_sgrna 30 --min_average_read_quality 30*).^36^ All described indel frequencies for each locus are determined by indel frequency in experimental sample minus the background rates in mock control sample.

### Cas12a RNP mediated knockout in NK cells

NK cells from PBMCs were isolated with EasySep™ Human NK Cell Isolation Kit (STEMCELL, 17955). NK cells were cultured in NK MACS Medium (MACS,130-114-429) for 7 days before electroporation. For each electroporation, 200 pmole crRNA and 100 pmole Cas12a were mixed at room temperature for 20 min, 1 million NK cells were resuspended in 20 μl Amaxa 4D nucleofector master mix (82% P3 + 18% supplement 1) (Lonza, V4XP-3032), then mixed with Cas12a RNPs for electroporation using program CM137. The electroporated NK cells were characterized after culturing in MACS medium for 5 days.

### Flow cytometry of NK cell surface markers

Cells were first stained with Live and Dead violet viability kit (Invitrogen, L-34963). To detect surface molecules, cells were stained in MACS buffer with target-specific antibodies for 30 min at 4°C in the dark. To detect IFN-γ (Biolegend, 502528) or CD96 (Biolegend, 338406), cells were stimulated with the IL-12 (10 ng/ml), IL-15 (50 ng/ml) and IL-18 (50 ng/ml) for 16 hrs, or with PMA and ionomycin (eBioscience, 00-4970-03) for 3 hrs. In both cases, protein transport inhibitors (eBioscience, 00-4980-03) were present during the stimulation. Example of gating strategy for flow cytometry analysis is shown in Supplementary Fig. S4.

### Cell culture and electroporation of human CD34+ HSPCs

Human CD34^+^ HSPCs from mobilized peripheral blood of deidentified healthy donors were obtained from Fred Hutchinson Cancer Research Center, Seattle, Washington. CD34^+^ HSPCs were enriched using the Miltenyi CD34 Microbead kit (Miltenyi Biotec). CD34^+^ HSPCs were thawed and cultured into X-VIVO 15 (Lonza, 04–418Q) supplemented with 100 ng ml^−1^ human SCF, 100 ng ml^−1^ human thrombopoietin (TPO) and 100 ng ml^−1^ recombinant human Flt3-ligand (Flt3-L). HSPCs were electroporated with RNP 24 h after thawing. Electroporation was performed using Lonza 4D Nucleofector (V4XP-3032 for 20 μl Nucleocuvette Strips) as the manufacturer’s instructions. The RNP complex was prepared by mixing Cas9 or Cas12a protein and sgRNA or crRNA at a 1:2.5 molar ratio and incubated for 15 min at room temperature immediately before electroporation. 50K HSPCs resuspended in 20 μl P3 solution were mixed with RNP and transferred to a cuvette for electroporation with program EO-100. The electroporated cells were resuspended with X-VIVO medium with cytokines and transferred into erythroid differentiation medium (EDM) consisting of IMDM supplemented with 330 μg ml^−1^ holo-human transferrin, 10 μg ml^−1^ recombinant human insulin, 2 IU ml^−1^ heparin, 5% human solvent detergent pooled plasma AB, 3 IU ml^−^ erythropoietin, 1% L-glutamine, and 1% penicillin/streptomycin 24 h later for *in vitro* differentiation. During days 0–7 of culture, EDM was further supplemented with 10^−6^ M hydrocortisone (Sigma), 100 ng ml^−1^ human SCF, and 5 ng ml^−1^ human IL-3 (R&D) as EDM-1. During days 7–11 of culture, EDM was supplemented with 100 ng ml^−1^ human SCF only as EDM-2. During days 11–18 of culture, EDM had no additional supplements as EDM-3. Fetal hemoglobin (HbF) induction was assessed on day 18 of erythroid culture.

### Hemoglobin HPLC

Hemolysates were prepared from erythroid cells after 18 days of erythroid differentiation for *in vitro* differentiation experiments. Hemolysate reagent (Helena Laboratories, 5125) was used, and samples were analyzed with D-10 Hemoglobin Analyzer (Bio-Rad). HbA2 and HbE cannot be distinguished by this method.

### Tn5 tagmentation and library preparation for GUIDE-tag

Adaptor oligonucleotides were synthesized by IDT (Supplementary Table S4).^37,38^ Transposon assembly was done by incubating 158 μg Tn5 with 1.4 nmol annealed oligo (contains the full-length Illumina forward (i5) adapter, a sample barcode, and unique molecule identifier (UMI) at room temperature for 60 min.^37^

Libraries were constructed as previously described.^37^ Briefly, 200 ng of genomic DNA was incubated with 2 μL of assembled transposome at 55 °C for 7 min, and the product was cleaned up (20 μL) with a Zymo column (Zymo Research, #D4013). Tagmented DNA was used for the 1^st^ PCR using PlatinumTM SuperFi DNA polymerase (Thermo) with i5 primer and GUIDE-Tag specific primers (Supplementary Table S5). (i5+Insert_F [GUIDE-tag_F] and i5+Insert_R [GUIDE-tag_R]). The i7 index was added in the 2^nd^ PCR and the PCR product was cleaned up with AMPure XP SPRI beads (Beckman Coulter, 0.9X reaction volume). Completed libraries were quantified by 4200 Tapestation System (Agilent) and Qubit 4.0 (Thermo Fisher), pooled with equal moles and sequenced with 150 bp paired-end reads on an Illumina MiniSeq instrument.

### GUIDE-tag data analysis

The GUIDE-tag raw sequencing data pre-processing and analysis pipeline is available at https://github.com/locusliu/GUIDESeq-Preprocess_from_Demultiplexing_to_Analysis and as we previously reported.^37,39,40^ Briefly, it consists of the following steps:

i. Demultiplexing and UMI extraction. Raw BCL files were converted and demultiplexed using the appropriate i5 and i7 sequencing barcodes, allowing up to one mismatch in each barcode. Unique molecular identifiers (UMIs) for each read were extracted for further downstream analysis.
ii. Raw reads were processed with fastqc (Version 0.11.9) and trim_galore (Version 0.6.5) (https://www.bioinformatics.babraham.ac.uk/projects/) to remove reads with low quality and trim adapters.
iii. Alignment analysis. Paired reads were then globally aligned (end-to-end mode) to human genome (hg38) and all the reference amplicons using bowtie2’s very sensitive parameter. Finally, Samtools (version 0.1.19) was used to create an index-sorted bam file.
iv. GUIDE-Tag sequencing data were run through the Bioconductor GUIDE-seq analysis pipeline (https://github.com/umasstr/GS-Preprocess). Briefly, for GUIDE-seq analysis processed paired reads were merged (if they overlap) and then globally aligned to the human genome (hg38) using bowtie2. Then BAM files and UMI files were used to aggregate unique reads. Mapped BAM files, sgRNA/crRNA fasta files and presorted UMI fastq files as inputs to run the Bioconductor GUIDE-Seq package to define potential off-target site across the whole genome. For off target site identification within potential peaks required the presence of a near-cognate recognition sequence for Cas9/Cas12a with these parameters: the maximum number of mismatches is 6 positions with one DNA/RNA bulge allowed and the presence of an NNG/NGN for SpyCas9 PAM or 5’ NNNN PAM for enAspCas12a.

### Statistical analyses

Statistical analysis for all transformed cell line experiments and CD34+ HSPC proof-of-concept experiments were performed with GraphPad Prism v8.4 (GraphPad Software, Inc., San Diego, CA). Statistical significance is determined by one-way or two-way ANOVA followed by multiple comparisons: *ns*, P > 0.05; *, P < 0.01; **, P < 0.001; ***, P < 0.0001; ****, P < 0.00001. R, a system for statistical computation and graphics, was used for the analysis of NK indel frequency and %Knockout, the CD34+ HSPC HF1 titration experiment, and CD34+ off-target amplicon-seq.^41^ Percentage of indel frequency, hemoglobin fraction, and knockout rate were first arcsine transformed to homogenize the variance. When Levene’s test indicates that the assumption of homogeneity of variances was met, one-way (Amplicon-seq) or two-way analysis of variance (ANOVA) with Randomized Complete Block Design was performed to test whether there are main effects of Site/Dose, Treatment, and whether there is a Dose/Site dependent Treatment effect. When there is no significant dose/site dependent treatment effect, the main effects of treatment and dose/site were reported. Otherwise, treatments were compared within each level of dose/site under the ANOVA framework using lsmeans package.^42^ P values were adjusted using the Hochberg method to correct for multiple inferences.^43^ When the assumption of homogeneity of variances was not met, unequal variance t-test was performed. All data is from three to six independent biological replicates. Error bars indicate ± S.E.M.

## Results

### Optimization of Cas12a nuclear localization signal sequence architecture

We previously reported achieving improvements in the editing efficiency for Cas12a and Cas9 in mammalian cells, primary cells, and zebrafish via modification of NLS sequence composition and architecture. However, there is still a gap in overall editing activity for Cas12a compared to Cas9 in primary CD34+ HSPCs.^14^ To further improve the efficiency of Cas12a targeted mutagenesis in primary cells, we applied principles that were successful for the 3xNLS SpyCas9 protein to AspCas12a, enAspCas12a, and LbaCas12a.^32^ We hypothesized that substitution of the SV40 NLS sequence with a more efficient nuclear import sequence, such as the c-Myc NLS, and an increase in the number of NLS sequences from two to three for our previously described 2xNLS-SV40-NLP Cas12a nuclease would facilitate more effective nuclear entry and result in a more robust genome editing platform.^44,45^

In our studies, we tested out new NLS designs directly as RNPs instead of plasmid encoded nucleases. Direct delivery of nucleases as RNPs into cells is an ideal vehicle for therapeutic applications due to their high editing activity.^46–48^ Furthermore, the short lifetime of RNPs in cells reduces the potential for off-target activity, which is a critical consideration for therapeutic genome editing approaches.^46–48^ Cas12a proteins with several different NLS sequence configurations were purified from an *E. Coli* overexpression system by Ni-NTA resin followed by cation exchange chromatography for comparative editing analysis (Fig. 1A). We first examined the influence of the different NLS frameworks on the editing activity of AspCas12a, enAspCas12a, and LbaCas12a at two previously characterized target sites (*AAVS1* site 1 and *EMX1* site 1) with RNPs delivered by electroporation to HEK293T cells at 5 pmol Cas12a protein:crRNA complex.^1,13,14^ At both genomic target sites, we observed that 2xNLS Cas12a with the substitution of SV40 NLS for the c-Myc NLS significantly increased the lesion frequency of the Cas12a nucleases by ~1.5-to-2-fold (Supplementary Fig. S1A; Supplementary Fig. S1B). Further significant improvement in editing activity was achieved by the addition of a third NLS (c-Myc NLS) to the C-terminus of 2xNLS-NLP-cMyc Cas12a in all the orthologs tested (Supplementary Fig. S1A; Supplementary Fig. S1B) Together, these results suggest that any Cas12a ortholog bearing three C-terminal NLS sequences, herein referred to as 3xNLS-NLP-cMyc-cMyc Cas12a, enables highly efficient gene editing in HEK293T cells.

To examine if these observations are general to a range of cell types, we used a more rigorously purified representative group of Cas12a variants (2xNLS-SV40-NLP AspCas12a, 2xNLS-NLP-cMyc AspCas12a, 3xNLS-NLP-cMyc-cMyc AspCas12a, 3xNLS-NLP-cMyc-cMyc enAspCas12a, 2xNLS-SV40-NLP LbaCas12a, 2xNLS-NLP-cMyc LbaCas12a, and 3xNLS-NLP-cMyc-cMyc LbaCas12a) to assess editing activity of the Cas12a RNPs at three genomic target sites (AAVS1 site 1 [Fig. 1B, 1C, and 1D], *EMX1* site 1 [Supplementary Fig. S1C], and *DNMT1* site 3 [Supplementary Fig. S1D]) in HEK293T, Jurkat, and K562 cells.^1,13^ Consistent with our previous observations, we found that the substitution of SV40 NLS for c-Myc NLS and the addition of a third C-terminal NLS resulted in a 1.25-to-3 fold increase in activity across all three cell lines (Fig. 1B, 1C, and 1D). Notably, the editing activity of 3xNLS-NLP-cMyc-cMyc AspCas12a and LbaCas12a were comparable to all versions of enAspCas12a, which is one of the most active Cas12a nucleases to date. The observation of improved activity by Cas12a regardless of target cell line suggests that our optimization to the NLS framework of Cas12a was applicable to a range of cell types. Based on its improved editing activity, we examined editing activity of Cas12a proteins with the 3xNLS-NLP-cMyc-cMyc (simplified as 3xNLS hereafter) framework in primary cells.

### Genome editing with new Cas12a NLS variants in Natural Killer cells

Natural killer (NK) cells are a subset of innate lymphoid cells (ILC) that are responsible for granzyme and perforin-mediated cytolytic activity against tumor and virus-infected cells.^49^ While NK cells are a well-studied subset of ILCs and possess promising clinical implications, reverse genetic screens to study gene function and the development of NK cell based therapeutics have lagged behind, likely due to challenges of genetic engineering NK cells using conventional methods. For example, retroviral transduction to deliver nucleases and shRNAs typically require a high viral titer and lentiviral transduction can be inconsistent for NK cells.^50^ Furthermore, plasmid expression of genome engineering tools have been limited in efficiency.^51^ Additionally, methods focused on delivering nucleases via mRNA require activation of NK cells prior to electroporation to make them more amenable to nucleic acid delivery.^52^ For these reasons, alternative strategies are being investigated, specifically those that employ electroporation of nucleases as RNPs to NK cells. While there have been some successes in genetically engineering NK cells with Cas9, many studies have been limited by lower indel frequencies (<80%), low knockout efficiency, or the requirement of activation of NK cells prior to electroporation.^50,53–55^ While editing with Cas9 has been suboptimal, alternative approaches utilizing Cas12a to engineer NK cells are being actively investigated.^29,51^ Based on our observation of improved editing by our new NLS framework in transformed cell lines and the intrinsic advantages of Cas12a listed above, we hypothesized that our new Cas12a NLS variants may serve as a more robust genome editing platform for NK cells.

To test our hypothesis, we delivered 1xNLS-NLP and the 3xNLS enAspCas12a targeting either the *IFNG* or *CD96* locus via electroporation into NK cells with two doses of RNP (50 pmol and 200 pmol) (Fig. 2A). At the *IFNG* target site, we found that the mean indel frequencies for 3xNLS enAspCas12a (71.16% and 84.61%) were significantly elevated compared to 1xNLS-NLP enAspCas12a (56.90% and 78.05%) (Fig. 2B; Supplementary Fig. S2A; Supplementary Fig. S2B). Additionally, comparison between 1xNLS and 3xNLS enAspCas12a at the IFNG target site, showed no discernable differences between the two NLS variants in both indel spectrum and indel distribution (Supplementary Fig. S2C; Supplementary Fig. S2D). Importantly, we observed only modest cellular toxicity associated with Cas12a treatment (Supplementary Fig. S2E). At the *CD96* target site, we found that the mean indel frequencies for 3xNLS enAspCas12a (73.87% and 86.96%) were significantly elevated compared to 1xNLS-NLP enAspCas12a (66.83% and 80.31%) (Fig. 2C; Supplementary Fig. S3A; Supplementary Fig. S3B). Like the IFNG target site, comparison between 1xNLS and 3xNLS enAspCas12a at the CD96 target site, showed no discernable differences between the two NLS variants in both indel spectrum and indel distribution (Supplementary Fig. S3C; Supplementary Fig. S3D). For both NLS variants at *IFNG* and *CD96*, approximately two-thirds (2/3) of the indels resulted in out-of-frame mutations (Supplementary Fig. S2D; Supplementary Fig. S3D). These observations suggest that the optimization of NLS architecture for Cas12a resulted in a nuclease with improved activity in NK cells, but overall similar properties with regards to editing outcomes.

**FIG 2.**
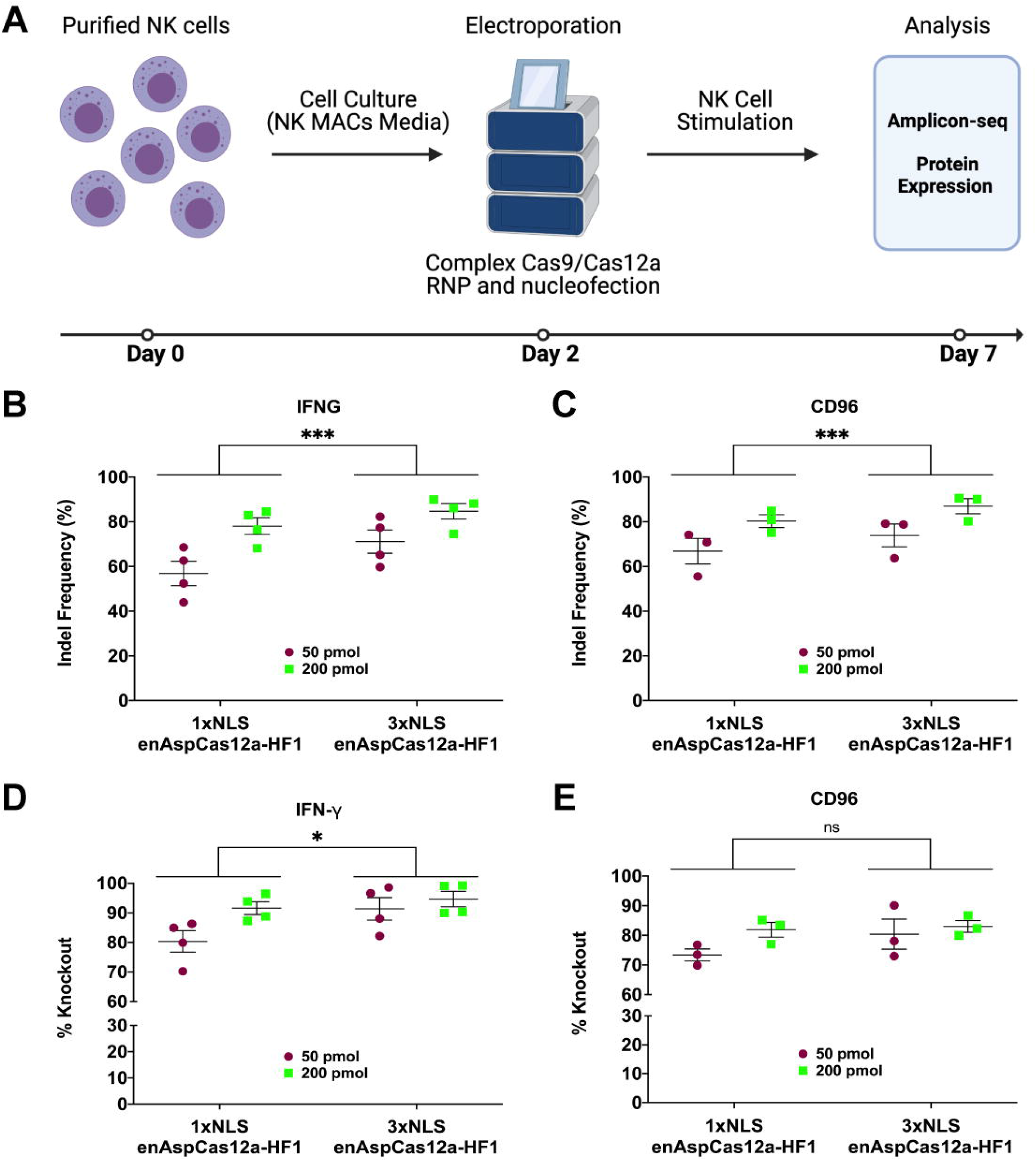
Robust gene editing in NK cells by enAspCas12a. (A) Schematic representation of experimental workflow for NK editing and phenotypic analysis. (B) Quantification of nuclease activity at *IFNG* (5 days post-nucleofection) based on Illumina sequencing. (C) Quantification of nuclease activity at *CD96* (5 days post-nucleofection) based on Illumina sequencing. (D) Quantification of %Knockout IFN-γ in stimulated NK cells. Cells are stimulated for 3 hours (PMA+ionomycin) or overnight (IL-12+IL-15+IL-18 cocktail). The fraction of cells positive for protein expression was determined by flow cytometry. (E) Quantification of %Knockout of CD96 in stimulated NK cells. Cells are stimulated for 3 hours (PMA+ionomycin) or overnight (IL-12+IL-15+IL-18 cocktail). The fraction of cells positive for protein expression was determined by flow cytometry. Results were obtained from four (IFNG) and three (CD96) independent experiments and presented as mean◻±◻SEM. *\*P*◻<◻0.05, *\*\*P*◻<◻0.01, \**\*\*P*◻<◻0.001 by two-way ANOVA with Tukey’s multiple comparisons test between 1xNLS and 3xNLS enAspCas12a.

Importantly, the impact of editing at each target site leads to robust phenotypic change. IFN-γ protein expression in NK cells can be stimulated through PMA + ionomycin or a IL-12+IL-15+IL-18 cytokine cocktail treatment.^51^ In stimulated cells, we found that both 1xNLS-NLP and 3xNLS enAspCas12a RNP efficiently disrupted the expression of IFN-γ (%Knockout > 80%) (Fig. 2D; Supplementary Fig. S2B). Comparing the %Knockout by 1xNLS-NLP and 3xNLS enAspCas12a, we observed that %Knockout of IFN-γ by 3xNLS enAspCas12a was significantly increased. Furthermore, we found that both 1xNLS-NLP and 3xNLS enAspCas12a RNP efficiently disrupted the expression of CD96 (%Knockout > 70%) (Fig. 2E; Supplementary Fig. S3B). However, we did not observe a significant difference between 1xNLS-NLP and 3xNLS enAspCas12a in %Knockout of CD96, likely due to the expression of CD96 in only a minor fraction of the NK cell population. These results demonstrate that enAspCas12a is a robust nuclease in NK cells, and that our new NLS architecture improves the editing activity and %Knockout efficiency of enAspCas12a in NK cells.

### 3xNLS enAspCas12a efficiently edits genetic regulatory elements in CD34+ HSPCs

To apply our new Cas12a nuclease platforms at a therapeutically relevant genomic locus, we tested the ability of enAspCas12a to disrupt the ATF4-binding motif within the *BCL11A* erythroid-lineage-specific +55 kb enhancer (Fig. 3A).^56,57^ Two previous studies revealed that ATF4 serves as an important regulator of globin expression.^56,57^ Depletion of ATF4 or disruption of the ATF4-binding motif in the +55kb enhancer in CD34+ HSPCs silences *BCL11A* expression in the erythroid lineage, resulting in increased production of fetal γ-globin protein in differentiated red blood cells.^56,57^ A similar induction of fetal γ-globin has been observed with the disruption of the GATA1-binding motif within the *BCL11A* erythroid-lineage-specific +58 kb enhancer in CD34+ HSPCs, which is currently under clinical development as an autologous bone marrow transplantation approach for the treatment of β-hemoglobinopathies.^32,58,59^ We examined the potency of HbF induction in the erythroid lineage when the ATF4 site within the *BCL11A* +55 kb enhancer is disrupted in CD34+ HSPCs.

**FIG 3.**
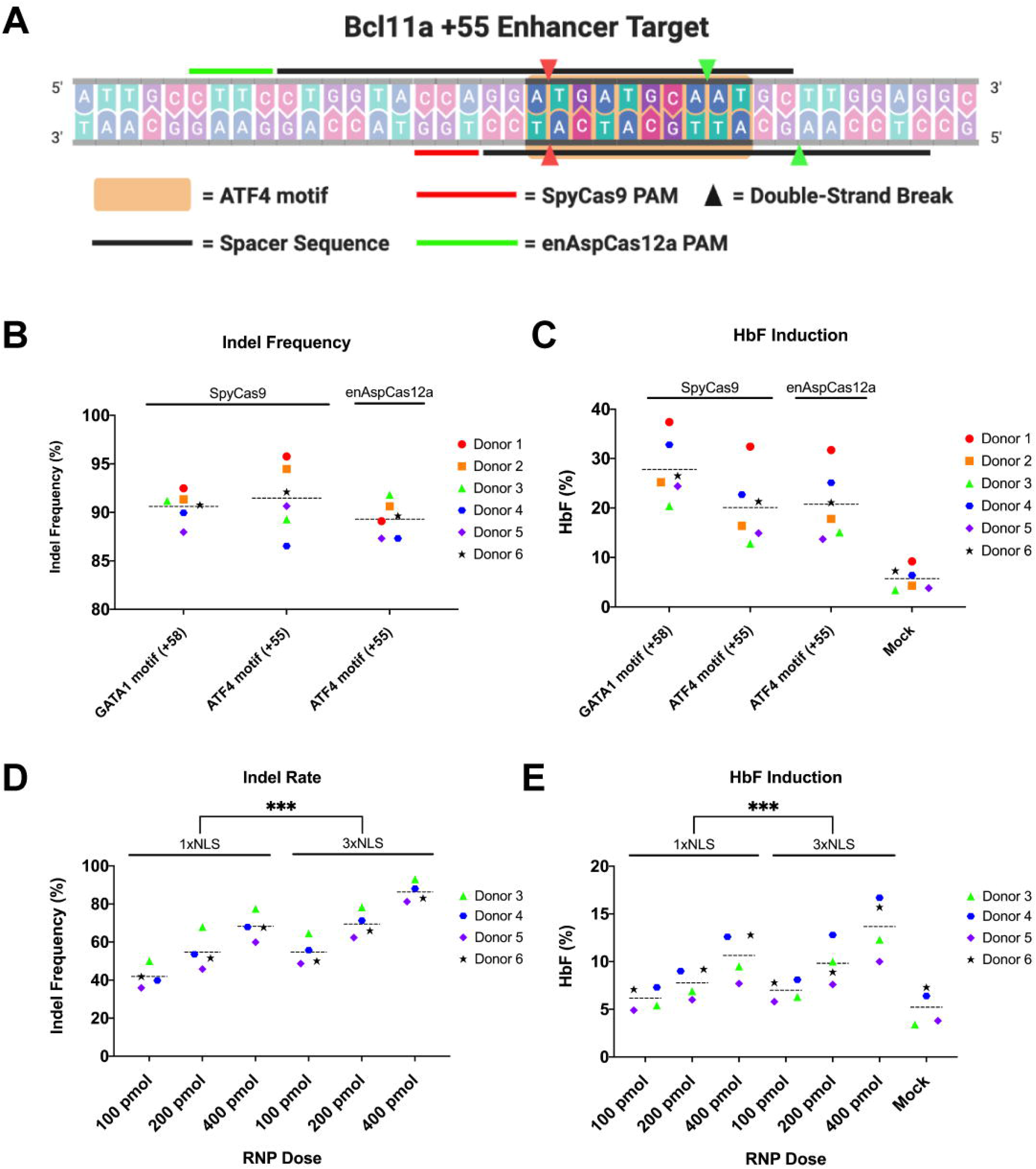
Disruption of ATF4-binding motif in +55 enhancer of *BCL11A* is an attractive alternative therapeutic approach. (A) Schematic representation of ATF4-binding motif in the +55 enhancer of *BCL11A* with indication of SpyCas9 and enAspCas12a target sites. (B) Quantification of editing efficiency at 3 days post-electroporation by SpyCas9 RNPs programmed with Spy1617 or SpyTS1 sgRNA, or enAspCas12a programmed with EnAspTS2 crRNA in CD34+ HSPCs from six donors. (C) Quantification of HbF induction in differentiated erythrocytes (18 days post-electroporation) CD34+ HSPCs from six donors treated with SpyCas9 RNPs programmed with Spy1617 or SpyTS1, or enAspCas12a programmed with EnAspTS2 crRNA. (D) Quantification of editing efficiency at 3 days post-electroporation by 1xNLS enAspCas12a-HF1 and 3xNLS enAspCas12a-HF1 at EnAspTS2 in CD34+ HSPCs from four donors. (E) Quantification of HbF induction in differentiated erythrocytes (18 days post-electroporation) disrupted in CD34+ HSPCs from four donors with 1xNLS enAspCas12a-HF1 and 3xNLS enAspCas12a-HF1 at EnAspTS2. HF1 denotes the high-fidelity version of the Cas12a nuclease. Each donor is represented by a specific-colored shape as noted in the legend. Dashed lines indicated the grand mean of the group. Results were obtained from six and four independent experiments and presented as mean◻±◻SEM. *ns* >◻0.05, *\*P*◻<◻0.05, *\*\*P*◻<◻0.01, \**\*\*P*◻<◻0.001 by two-way ANOVA with Tukey’s multiple comparisons test between 1xNLS and 3xNLS enAspCas12a.

As a proof-of-concept experiment, we utilized 3xNLS enAspCas12a to target the ATF4 site in the +55 enhancer of *BCL11A*. enAspCas12a can recognize more diverse PAM sequences than AspCas12a and LbaCas12a, which allows enAspCas12a to be positioned for DNA cleavage within the ATF4 target sequence (EnAspTS2; Fig. 3A). To provide a comparison to an orthogonal nuclease, we examined editing efficiency at a previously validated SpyCas9 target site overlapping the ATF4-binding motif (SpyTS1; Fig. 3A).^56^ We performed pilot experiments with the active RNP complexes in CD34+ HSPCs from six healthy donors. CD34+ HSPCs were electroporated with 200 pmol SpyCas9 or enAspCas12a complexed with their respective guide RNAs targeting the ATF4-binding motif in the +55 enhancer of *BCL11A* (SpyTS1 and enAspTS2). Editing by 200 pmol SpyCas9 targeting the GATA1-binding motif in the +58 enhancer of *BCL11A* (Spy1617) served as a benchmark and positive control for HbF induction.^32,33,60^ We found that all SpyCas9 and enAsCas12a RNPs were able to effectively edit their target sites (indel frequencies >85%) (Fig. 3B). Importantly, the mean editing rates for both Cas9 and Cas12a were approximately 90% (Spy1617 – 90.61%, SpyTS1 – 91.56%, and EnAspTS2 – 89.30%). Interestingly, we observed a notable difference in the indel spectrums produced by the sgRNA for SpyTS1 and crRNA for EnAspTS2 (Supplementary Fig. S5A). With EnAspTS2, most of the products had larger deletions (≥ 5bp) within the ATF4-binding motif. Comparatively, for the SpyTS1 sgRNA, we observed that most of the products had smaller insertions or deletions that were on the distal 3’ end of the ATF4-binding motif. Furthermore, indel distribution analysis revealed that Cas12a produced a higher percentage of larger deletions (≥ 5bp) (88.6% of all mapped reads, than Cas9 (32.54% of all mapped reads) at the ATF4 target site (Supplementary Fig. S5B). In addition to examining editing activity, we were interested in the potential effects on HbF induction in differentiated erythrocytes. Importantly, editing of *BCL11A* target sites led to robust HbF protein expression in erythroid progenitor cells differentiated from the edited CD34+ HSPCs (Fig. 3C). Consistent with data from previous reports, HbF induction rates for erythroid progenitors derived from edited CD34+ HSPCs are the highest for the Spy1617 target site in the +58 enhancer compared to mock treated cells (27.78%, comparison to mock p-value <0.0001).^32,59^ Modestly lower HbF induction levels are achieved for editing in the ATF4 site by SpyTS1 (20.08%, comparison to mock p-value <0.0009) and EnAspTS2 (20.75%, comparison to mock p-value <0.0005).

Given the promising results with 3xNLS enAspCas12a editing at the ATF4-binding motif, we examined the efficacy of the high-fidelity version of enAspCas12 (HF1) since this protein displays dramatically decreased off-target editing rates.^25^ We performed a titration experiment with 1xNLS-NLP and 3xNLS enAspCas12a-HF1 RNPs targeting the ATF4 TS2 in CD34+ HSPCs with four different donors. We observed dose-dependent editing for 1xNLS-NLP and 3xNLS enAspCas12a-HF1 (Fig. 3D; Supplementary Fig. S5C). Furthermore, the overall editing activity of the new 3xNLS format was significantly improved compared to 1xNLS-NLP, regardless of RNP amount. Additionally, HbF levels in erythroid progenitors were significantly elevated for 3xNLS enAspCas12a-HF1 relative to the 1xNLS-NLP (Fig. 3E; Supplementary Fig. S5D). Interestingly, the HbF induction levels of the HF1 variants were lower relative to the wild-type variant of enAspCas12a, likely due to the lower overall editing activity of the HF1 variant. Together, these results suggest that the new Cas12a NLS variants are a promising platform for the disruption of functional sequence motifs in therapeutically relevant genomic loci and that the 3xNLS architecture may improve the activity of other Cas12a orthologs for editing in primary hematopoietic cells.

### Characterization of Cas12a specificity with new NLS frameworks

While the improvement in activity is critical for the use of Cas12a for research and therapeutic applications, it is important to verify that the new NLS architectures do not decrease the high intrinsic specificity of Cas12a. To compare the specificity of the Cas12a proteins with different NLS frameworks, we used a crRNA targeting *DNMT1* site 3, which has a well-characterized off-target profile by both targeted deep sequencing and unbiased genome-wide off-target detection approaches (Fig. 4A).^9,10,13^ We first carried out an initial titration experiment in HEK293T cells to determine the optimal amount of RNP where all Cas12a variants would have similar near-saturating on-target editing activity. We observed that the most active variants (3xNLS AspCas12a and 3xNLS enAspCas12a) required delivery of lower amounts of Cas12a protein-crRNA complex to achieve similar on-target nuclease activities (Supplementary Fig. S6A). Next, we nucleofected HEK293T cells with doses of each RNP targeting *DNMT1* site 3 to achieve ~80% editing and performed targeted deep sequencing analyses of PCR products spanning the target site and 13 previously identified off-target sites. Comparison of our previously described 2xNLS-SV40-NLP Cas12a with our new Cas12a NLS variants (2xNLS-NLP-cMyc Cas12a and 3xNLS-NLP-cMyc-cMyc Cas12a) demonstrates that neither the substitution of the SV40 NLS for c-Myc NLS nor the addition of a third NLS sequence significantly increase the lesion frequency at any of the previously validated off-target sites (Fig. 4A). Furthermore, enAspCas12a was the most promiscuous nuclease, with substantially more editing observed at all 13 off-target sites, consistent with previous examinations of the specificity of enAspCas12a (Fig. 4A).^25^

**FIG 4.**
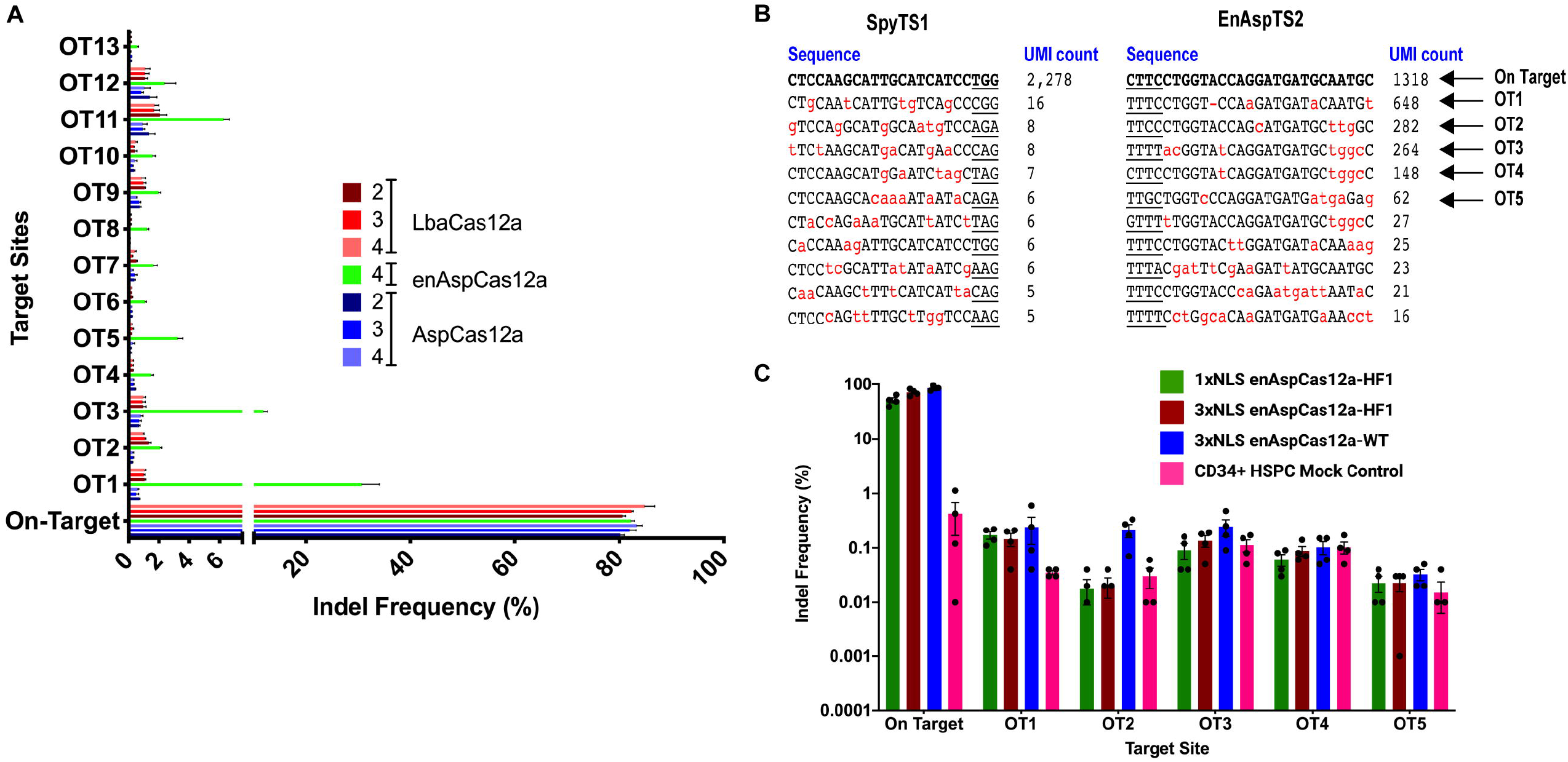
Cas12a NLS variants do not demonstrate increased activity at off-target sites. (A) Quantification of editing activity of various Cas12a nucleases programmed with the crRNA targeting *DNMT1* site 3 at the on-target and 13 GUIDE-seq and computationally identified potential off-target sites. The Cas12a proteins were delivered at specific amounts of protein:crRNA complex (2xNLS-NLP-SV40 AspCas12a: 5 pmol; 2xNLS-NLP-cMyc AspCas12a: 5 pmol; 3xNLS-NLP-cMyc-cMyc AspCas12a: 2.5 pmol; 3xNLS-NLP-cMyc-cMyc enAspCas12a: 1.25 pmol; 2xNLS-NLP-SV40 LbaCas12a: 40 pmol; 2xNLS-NLP-cMyc LbaCas12a: 40 pmol; 3xNLS-NLP-cMyc-cMyc Cas12a: 40 pmol) to achieve approximately 80% editing at the target site. Statistical analysis between the NLS variants were determined by two-way Anova with multiple comparisons. (B) GUIDE-tag analysis of SpyCas9 programmed with SpyTS1 sgRNA or enAspCas12a programmed with EnAspTS2 crRNA. Sequences shown are the top 10 recovered sites based on UMI count. Underline indicates the PAM of the nuclease. Red letters indicate mismatches with the cognate spacer sequence. Arrows indicate off-target (OT) sites further tested in CD34+ HSPCs. (C) Quantification of editing activity by enAspCas12a variants programmed with EnAspTS2 crRNA at on-target and top 5 GUIDE-tag identified potential off-target sites in CD34+ HSPCs. Results were obtained from three (*DNMT1* site 3) and four (enAspTS2) independent experiments and presented as mean◻±◻SEM.

After testing the specificity of 3xNLS Cas12a framework at a previously validated site in HEK293T cells, we evaluated the specificity of our improved Cas12a NLS variants using guides designed for the disruption of ATF4-binding motif in the *BCL11A* +55 enhancer. We assayed for active off-target sites for 3xNLS enAspCas12a and SpyCas9 with corresponding guide RNAs by GUIDE-tag genome-wide specificity analysis in HEK293T cells.^37^ A small number of potential off-target sites were identified for both SpyTS1 and enAspTS2 target sites as determined by Unique Molecular Identifier (UMI) count in GUIDE-tag analysis (Fig. 4B; Supplementary Table S6 [SpyTS1]; Supplementary Table S7 [EnAspTS2]).

To assess the specificity of our 3xNLS enAspCas12a variants in primary cells, we performed amplicon deep sequencing at the top five potential off-target sites identified from the GUIDE-tag analysis in Cas12a treated CD34+ HSPCs and HEK293T cells. Consistent with the GUIDE-tag analysis, high levels of editing were observed at OT1 in addition to more modest levels of editing at OT2, OT3, OT4, and OT5 in HEK293T cells (Supplementary Fig. S6B). In CD34+ HSPCs, deep sequencing analysis indicates very low mutagenesis at the top five potential off-target sites in CD34+ HSPCs for all the enAspCas12a NLS variants tested (Fig. 4C). However, we saw higher editing activity relative to the mock control background at both, OT1 and OT2 for EnAspTS2 (Fig. 4C). Additionally, we observed indels in the sequences associated with the 1xNLS-HF1, 3xNLS-HF1, and 3xNLS-WT enAspCas12a variants at OT1 (Supplementary Fig. S6C; Supplementary Table S8) and 3xNLS-WT at OT2 (Supplementary Fig. S6C; Supplementary Table S9) that are consistent with Cas12a editing products. Together, these data suggest that there may be two minor off-target sites for enAspCas12a programmed with an EnAspTS2 crRNA.

## Discussion

Type V CRISPR-Cas12a nucleases are a well-characterized gene-editing platform that has been used for editing in vertebrate and invertebrate systems. Previous studies have shown that the type of NLS framework impacts the efficiency of genome editing by SpyCas9 and Cas12a in transformed mammalian cell lines, CD34+ HSPCs, and zebrafish.^13,14,18,21–23,25,27^ However, the efficiency of mutagenesis by Cas12a is typically lower than SpyCas9, especially in quiescent primary cells, therefore potentially limiting its utilization as a therapeutic agent.^14^

Here, we describe improvements to Cas12a nucleases that improve indel frequencies in transformed mammalian cell lines and therapeutically relevant primary cells via improved nuclear entry. Similar to previous reports described for SpyCas9, we found that the number and composition of the NLSs significantly impact the nuclease activity of Cas12a.^32^ More specifically, substitution of the previously utilized SV40 NLS for the more efficient, c-Myc NLS, and addition of a third NLS to the C-terminal of Cas12a resulted in a more robust genome editing platform. This enhancement of activity was observed in all Cas12a orthologs tested (Asp, enAsp, or Lba), which suggests that the 3xNLS framework can be utilized to further improve the activity of other Cas12a nucleases not explored in this study.^28–30^ Although we demonstrated that the c-Myc NLS is a superior import sequence for Cas12a than SV40 NLS, other classical and non-classical NLS sequences could be explored in the future to further optimize the Cas12a platform.^61^

We examined the impact of NLS architecture on editing activity in two relevant primary cell types, natural killer cells and CD34+ HSPCs. In NK cells, we compared the activity of the original 1xNLS-NLP and 3xNLS NLS architectures for enAspCas12a. We found the inclusion of optimization of the NLS framework significantly improved the indel frequencies of Cas12a in NK cells. Furthermore, we found the knockout of *IFNG* was significantly improved in our 3xNLS format compared to the previously reported 1xNLS-NLP framework. In a recent study by Zhang *et al*. demonstrated robust editing activity in NK cells using AspCas12a mutant referred as “AsCas12a Ultra” (M537R and F870L mutations).^29^ Similar to the study by Zhang *et al*., we demonstrated robust genome editing with Cas12a nucleases in NK cells.^29,51^ The ability to effectively mutagenize NK cells with Cas12a nuclease may prove valuable when considering reverse genetic screens for studying the function of NK cells or for therapeutic applications such as the generation of chimeric antigen receptor (CAR)-engineered NK (CAR-NK) cells.^62^

Likewise, we observed that optimization to Cas12a via modifications to the NLS architecture produced a more robust genome editing platform in CD34+ HSPCs. Specifically, comparison between 1xNLS-NLP and 3xNLS enAspCas12a-HF1 demonstrated that there was significant improvement in editing activity. Across four donors and three nuclease doses, editing CD34+ HSPCs with 3xNLS enAspCas12a-HF1 led to significantly more robust HbF induction compared to 1xNLS-NLP enAspCas12a-HF1 in differentiated erythrocytes. Based on the data for NK cells and CD34+ HSPCs, we believe that the optimized NLS composition and architecture described herein will improve Cas12a activity broadly across therapeutically relevant primary cell types.

Additionally, when we compared our 3xNLS enAspCas12a induced disruption of the ATF4-binding motif at the +55 enhancer of *BCL11A* strategy to previously validated and currently investigated strategies (Spy1617 and SpyTS1), we found that our Cas12a-based approach is comparable both in overall editing activity in CD34+ HSPCs and HbF induction in differentiated erythrocytes. Interestingly, despite the difference in indel products produced by SpyTS1 and EnAsp2, we did not observe a difference in HbF induction. This suggests that even minimal disruption of the ATF4-binding motif is sufficient to disrupt *BCL11A* expression, resulting in robust HbF induction. However, the observed editing outcomes at the ATF4 target site suggest that Cas12a may provide a superior nuclease to Cas9 for regulatory element disruption when larger deletions are required. Similar to a study by Huang *et al*., we observed that disruption of the ATF4-binding motif at the +55 enhancer of *BCL11A* was modestly less efficacious to the disruption of the GATA1-binding motif at the +58 enhancer of *BCL11A* for the induction of HbF in differentiated erythrocytes.^38^ However, compared to the study by Huang *et* al., we observed substantially higher induction of HbF.^56^ This is potentially due to higher editing activity of our Cas9 and Cas12a proteins in CD34+ HSPCs. Given the comparable editing activity and therapeutically relevant levels of HbF induced by Cas9 (SpyTS1) and Cas12a (EnAspTS2) via disruption of the ATF4-binding motif at the +55 enhancer of *BCL11A*, therapeutic strategies targeting this regulatory element could be particularly useful as an alternative or combinatorial therapeutic approach for the amelioration of β-hemoglobinopathies.

Lastly, we found that our Cas12a NLS variants did not demonstrate increased off-target activity in HEK293T cells at a commonly utilized target site (*DNMT1* site 3). Furthermore, we performed an unbiased genome-wide off-target identification method (GUIDE-tag) to detect potential near cognate sites for SpyTS1 and EnAspTS2 in HEK293T cells. To our knowledge this is the first study to examine the potential off-target activity of SpyTS1. Based on our preliminary examination of SpyTS1, it appears to be a highly specific target site. For enAspCas12a programmed with the EnAspTS2 crRNA, we did not see substantial editing activity at the top potential off-target sites in CD34+ HSPCs. However, the indel frequencies and indel spectrum produced by the NLS variants at OT1 and OT2 suggest that these off-target sites may be active at a low rate for the wild-type variant of 3xNLS enAspCas12a. Furthermore, with the high-fidelity variant of 3xNLS enAspCas12a, we saw that there was a decrease in observable editing at OT2. However, it appeared as though OT1 was still active despite the use of a high-fidelity nuclease. To eliminate the editing activity at these off-target sites, can examine potential protein or guide RNA specific modifications to Cas12a and its crRNA to improve specificity of wild-type variant of 3xNLS enAspCas12a.^25,28,63^ More analysis is needed to fully examine the potential genotoxic effects, such as large genomic rearrangements, deletions and chromothripsis, produced by targeting the ATF4-binding motif in the +55 enhancer of *BCL11A* with enAspCas12a.^64–71^

## Conclusions

We have demonstrated that optimization to the NLS framework of Cas12a is an effective method to improve the mutagenesis frequencies of the nucleases. Here, we demonstrate more robust genome editing with Cas12a in mammalian and primary cells, relative to previous Cas12a variants. We show that our NLS optimization approach can be applied to various Cas12a orthologs resulting in high editing activity - approaching ~90% in many cases - without sacrificing the high intrinsic specificity of Cas12a nucleases. Furthermore, we applied the NLS-optimized enAspCas12a to disrupt the ATF4-binding motif at the +55 enhancer of *BCL11A* as a potential therapeutic strategy for the amelioration of β-hemoglobinopathies. We show that this therapeutic approach is similar in HbF induction to current clinical strategies being investigated. Additionally, our genome-wide analysis of off-target activity suggests that the Cas9 and Cas12a target sites overlapping the ATF4-binding motif permit efficient editing of this locus with minimal off-target events, which is an important consideration for therapeutic application. In conclusion, our findings suggest that Cas12a systems with improved NLS architectures provide a platform that will have broad utility for both research and therapeutic applications.

## Acknowledgements

All new reagents described in this work are being deposited with the nonprofit plasmid-distribution service Addgene. We thank E. Sonthiemer, M. Socolovsky, C. Peterson, and M. Brodsky (University of Massachusetts Chan Medical School) for critical input of the manuscript.

## Author Contributions

K.L., P.P.L., J.Z., D.E.B. and S.A.W. designed the experiments. K.L. purified proteins and performed experiments in transformed cell lines with assistance from P.P.L., S.A.M., F.I., K.P., and S.A.W.. Y.W. isolated and performed experiments in NK cells with assistance from K.L.. J.Z. performed experiments in CD34+ HSPCs with assistance from K.L.. K.L., P.P.L., L.J.Z., and S.A.W. analyzed the experimental data. K.L. and S.A.W. wrote the manuscript, which was revised and approved by all authors.

## References

1. Zetsche B, Gootenberg JS, Abudayyeh OO, et al. Cpf1 is a single RNA-guided endonuclease of a class 2 CRISPR-Cas system. Cell 2015;163:759–771.

2. Shmakov S, Abudayyeh OO, Makarova KS, et al. Discovery and Functional Characterization of Diverse Class 2 CRISPR-Cas Systems. Mol Cell 2015;60:385–397.

3. Fagerlund RD, Staals RHJ, Fineran PC. The Cpf1 CRISPR-Cas protein expands genome-editing tools. Genome Biol 2015;16:251.

4. Kim HK, Song M, Lee J, et al. In vivo high-throughput profiling of CRISPR–Cpf1 activity. Nat Methods. Epub ahead of print 2016. DOI: 10.1038/nmeth.4104.

5. Fonfara I, Richter H, Bratovič M, et al. The CRISPR-associated DNA-cleaving enzyme Cpf1 also processes precursor CRISPR RNA. Nature 2016;1–19.

6. Port F, Starostecka M, Boutros M. Multiplexed conditional genome editing with Cas12a in Drosophila. Proc Natl Acad Sci U S A 2020;117:22890–22899.

7. Campa CC, Weisbach NR, Santinha AJ, et al. Multiplexed genome engineering by Cas12a and CRISPR arrays encoded on single transcripts. Nat Methods 2019;16:887–893.

8. Weisbach NR, Meijs A, Platt RJ. Multiplexed Genome Engineering with Cas12a. Methods Mol Biol 2021;2312:171–192.

9. Kim D, Kim J, Hur JK, et al. Genome-wide analysis reveals specificities of Cpf1 endonucleases in human cells. Nat Biotechnol 2016;34:863–868.

10. Kleinstiver BP, Tsai SQ, Prew MS, et al. Genome-wide specificities of CRISPR-Cas Cpf1 nucleases in human cells. Nat Biotechnol 2016;34:869–874.

11. Yan WX, Mirzazadeh R, Garnerone S, et al. BLISS is a versatile and quantitative method for genome-wide profiling of DNA double-strand breaks. Nat Commun 2017;8:15058.

12. Port F, Bullock SL. Augmenting CRISPR applications in Drosophila with tRNA-flanked sgRNAs. Nat Methods 2016;13:852–854.

13. Liu P, Luk K, Shin M, et al. Enhanced Cas12a editing in mammalian cells and zebrafish. Nucleic Acids Res 2019;47:4169–4180.

14. Xu S, Luk K, Yao Q, et al. Editing aberrant splice sites efficiently restores β-globin expression in β-thalassemia. Blood. Epub ahead of print January 31, 2019. DOI: 10.1182/blood-2019-01-895094.

15. Hur JK, Kim K, Been KW, et al. Targeted mutagenesis in mice by electroporation of Cpf1 ribonucleoproteins. Nat Biotechnol 2016;34:807–808.

16. Watkins-Chow DE, Varshney GK, Garrett LJ, et al. Highly Efficient Cpf1-Mediated Gene Targeting in Mice Following High Concentration Pronuclear Injection. G3 2017;7:719–722.

17. Kim Y, Cheong S-A, Lee JG, et al. Generation of knockout mice by Cpf1-mediated gene targeting. Nat Biotechnol 2016;34:808–810.

18. Dumeau C-E, Monfort A, Kissling L, et al. Introducing gene deletions by mouse zygote electroporation of Cas12a/Cpf1. Transgenic Res 2019;28:525–535.

19. Ahn W-C, Park K-H, Bak IS, et al. In vivo genome editing using the Cpf1 ortholog derived from Eubacterium eligens. Sci Rep 2019;9:13911.

20. Moreno-Mateos MA, Fernandez JP, Rouet R, et al. CRISPR-Cpf1 mediates efficient homology-directed repair and temperature-controlled genome editing. Nat Commun 2017;8:2024.

21. Bernabé-Orts JM, Casas-Rodrigo I, Minguet EG, et al. Assessment of Cas12a-mediated gene editing efficiency in plants. Plant Biotechnol J 2019;17:1971–1984.

22. Banakar R, Schubert M, Collingwood M, et al. Comparison of CRISPR-Cas9/Cas12a Ribonucleoprotein Complexes for Genome Editing Efficiency in the Rice Phytoene Desaturase (OsPDS) Gene. Rice 2020;13:4.

23. Li B, Rui H, Li Y, et al. Robust CRISPR/Cpf1 (Cas12a)-mediated genome editing in allotetraploid cotton (Gossypium hirsutum). Plant Biotechnol J 2019;17:1862–1864.

24. Gao L, Cox DBT, Yan WX, et al. Engineered Cpf1 variants with altered PAM specificities. Nat Biotechnol 2017;35:789–792.

25. Kleinstiver BP, Sousa AA, Walton RT, et al. Engineered CRISPR-Cas12a variants with increased activities and improved targeting ranges for gene, epigenetic and base editing. Nat Biotechnol 2019;37:276–282.

26. Teng F, Li J, Cui T, et al. Enhanced mammalian genome editing by new Cas12a orthologs with optimized crRNA scaffolds. Genome Biol 2019;20:15.

27. Tóth E, Varga É, Kulcsár PI, et al. Improved LbCas12a variants with altered PAM specificities further broaden the genome targeting range of Cas12a nucleases. Nucleic Acids Res 2020;48:3722–3733.

28. Zhou J, Chen P, Wang H, et al. Cas12a variants designed for lowergenome-wide off-target effectthrough stringent PAM recognition. Mol Ther. Epub ahead of print October 20, 2021. DOI: 10.1016/j.ymthe.2021.10.010.

29. Zhang L, Zuris JA, Viswanathan R, et al. AsCas12a ultra nuclease facilitates the rapid generation of therapeutic cell medicines. Nat Commun 2021;12:3908.

30. Zhu D, Wang J, Yang D, et al. High-Throughput Profiling of Cas12a Orthologues and Engineered Variants for Enhanced Genome Editing Activity. Int J Mol Sci;22. Epub ahead of print December 10, 2021. DOI: 10.3390/ijms222413301.

31. Tran MH, Park H, Nobles CL, et al. A more efficient CRISPR-Cas12a variant derived from Lachnospiraceae bacterium MA2020. Molecular Therapy - Nucleic Acids 2021;24:40–53.

32. Wu Y, Zeng J, Roscoe BP, et al. Highly efficient therapeutic gene editing of human hematopoietic stem cells. Nat Med. Epub ahead of print March 25, 2019. DOI: 10.1038/s41591-019-0401-y.

33. Demirci S, Zeng J, Wu Y, et al. BCL11A enhancer-edited hematopoietic stem cells persist in rhesus monkeys without toxicity. J Clin Invest 2020;130:6677–6687.

34. Iyer S, Mir A, Vega-Badillo J, et al. Efficient Homology-directed Repair with Circular ssDNA Donors. bioRxiv 2019;864199.

35. Bolukbasi MF, Gupta A, Oikemus S, et al. DNA-binding-domain fusions enhance the targeting range and precision of Cas9. Nat Methods 2015;12:1150–1156.

36. Clement K, Rees H, Canver MC, et al. CRISPResso2 provides accurate and rapid genome editing sequence analysis. Nat Biotechnol 2019;37:224–226.

37. Liang S-Q, Liu P, Smith JL, et al. Genome-wide detection of CRISPR editing in vivo using GUIDE-tag. Nat Commun 2022;13:1–14.

38. Giannoukos G, Ciulla DM, Marco E, et al. UDiTaS™, a genome editing detection method for indels and genome rearrangements. BMC Genomics 2018;19:212.

39. Rodríguez TC, Dadafarin S, Pratt HE, et al. Chapter Three - Genome-wide detection and analysis of CRISPR-Cas off-targets. In: Singh V (ed) Progress in Molecular Biology and Translational Science. Academic Press; pp. 31–43.

40. Zhu LJ, Lawrence M, Gupta A, et al. GUIDEseq: a bioconductor package to analyze GUIDE-Seq datasets for CRISPR-Cas nucleases. BMC Genomics 2017;18:379.

41. Ihaka R, Gentleman R. R: A Language for Data Analysis and Graphics. J Comput Graph Stat 1996;5:299–314.

42. Lenth RV. Least-Squares Means: The R Package lsmeans. J Stat Softw 2016;69:1–33.

43. Huang Y, Hsu JC. [No title]. Biometrika 2007;94:965–975.

44. Ray M, Tang R, Jiang Z, et al. Quantitative tracking of protein trafficking to the nucleus using cytosolic protein delivery by nanoparticle-stabilized nanocapsules. Bioconjug Chem 2015;26:1004–1007.

45. Dang CV, Lee WM. Identification of the human c-myc protein nuclear translocation signal. Mol Cell Biol 1988;8:4048–4054.

46. Moon SB, Kim DY, Ko J-H, et al. Recent advances in the CRISPR genome editing tool set. Exp Mol Med 2019;51:1–11.

47. Zhang S, Shen J, Li D, et al. Strategies in the delivery of Cas9 ribonucleoprotein for CRISPR/Cas9 genome editing. Theranostics 2021;11:614–648.

48. Chandrasekaran AP, Song M, Kim K-S, et al. Different Methods of Delivering CRISPR/Cas9 Into Cells. Prog Mol Biol Transl Sci 2018;159:157–176.

49. Abel AM, Yang C, Thakar MS, et al. Natural Killer Cells: Development, Maturation, and Clinical Utilization. Front Immunol 2018;9:1869.

50. Rautela J, Surgenor E, Huntington ND. Drug target validation in primary human natural killer cells using CRISPR RNP. J Leukoc Biol 2020;108:1397–1408.

51. Wang Y, Lifshitz L, Silverstein NJ, et al. Clarification of human blood ILC subtype interrelatedness and discovery of amphiregulin production by human NK cells shed light on HIV-1 pathogenesis. bioRxiv 2021;2021.04.20.440368.

52. Pomeroy EJ, Hunzeker JT, Kluesner MG, et al. A Genetically Engineered Primary Human Natural Killer Cell Platform for Cancer Immunotherapy. Mol Ther 2020;28:52–63.

53. Huang R-S, Lai M-C, Shih H-A, et al. A robust platform for expansion and genome editing of primary human natural killer cells. J Exp Med;218. Epub ahead of print March 1, 2021. DOI: 10.1084/jem.20201529.

54. Lambert M, Leijonhufvud C, Segerberg F, et al. CRISPR/Cas9-Based Gene Engineering of Human Natural Killer Cells: Protocols for Knockout and Readouts to Evaluate Their Efficacy. Methods Mol Biol 2020;2121:213–239.

55. Huang R-S, Shih H-A, Lai M-C, et al. Enhanced NK-92 Cytotoxicity by CRISPR Genome Engineering Using Cas9 Ribonucleoproteins. Front Immunol 2020;11:1008.

56. Huang P, Peslak SA, Lan X, et al. The HRI-regulated transcription factor ATF4 activates BCL11A transcription to silence fetal hemoglobin expression. Blood 2020;135:2121–2132.

57. Boontanrart MY, Schröder MS, Stehli GM, et al. ATF4 Regulates MYB to Increase γ-Globin in Response to Loss of β-Globin. Cell Rep 2020;32:107993.

58. Chang K-H, Smith SE, Sullivan T, et al. Long-Term Engraftment and Fetal Globin Induction upon BCL11A Gene Editing in Bone-Marrow-Derived CD34+ Hematopoietic Stem and Progenitor Cells. Mol Ther Methods Clin Dev 2017;4:137–148.

59. Frangoul H, Altshuler D, Cappellini MD, et al. CRISPR-Cas9 Gene Editing for Sickle Cell Disease and β-Thalassemia. N Engl J Med 2021;384:252–260.

60. Zeng J, Wu Y, Ren C, et al. Therapeutic base editing of human hematopoietic stem cells. Nat Med 2020;26:535–541.

61. Lu J, Wu T, Zhang B, et al. Types of nuclear localization signals and mechanisms of protein import into the nucleus. Cell Commun Signal 2021;19:60.

62. Xie G, Dong H, Liang Y, et al. CAR-NK cells: A promising cellular immunotherapy for cancer. EBioMedicine 2020;59:102975.

63. Kim H, Lee W-J, Oh Y, et al. Enhancement of target specificity of CRISPR-Cas12a by using a chimeric DNA-RNA guide. Nucleic Acids Res 2020;48:8601–8616.

64. Kim D, Luk K, Wolfe SA, et al. Evaluating and Enhancing Target Specificity of Gene-Editing Nucleases and Deaminases. Annu Rev Biochem. Epub ahead of print March 18, 2019. DOI: 10.1146/annurev-biochem-013118-111730.

65. Haapaniemi E, Botla S, Persson J, et al. CRISPR-Cas9 genome editing induces a p53-mediated DNA damage response. Nat Med. Epub ahead of print June 11, 2018. DOI: 10.1038/s41591-018-0049-z.

66. Ihry RJ, Worringer KA, Salick MR, et al. p53 inhibits CRISPR-Cas9 engineering in human pluripotent stem cells. Nat Med. Epub ahead of print June 11, 2018. DOI: 10.1038/s41591-018-0050-6.

67. van den Berg J, G Manjón A, Kielbassa K, et al. A limited number of double-strand DNA breaks is sufficient to delay cell cycle progression. Nucleic Acids Res 2018;46:10132–10144.

68. Shin HY, Wang C, Lee HK, et al. CRISPR/Cas9 targeting events cause complex deletions and insertions at 17 sites in the mouse genome. Nat Commun 2017;8:15464.

69. Adikusuma F, Piltz S, Corbett MA, et al. Large deletions induced by Cas9 cleavage. Nature 2018;560:E8–E9.

70. Kosicki M, Tomberg K, Bradley A. Repair of double-strand breaks induced by CRISPR-Cas9 leads to large deletions and complex rearrangements. Nat Biotechnol. Epub ahead of print July 16, 2018. DOI: 10.1038/nbt.4192.

71. Leibowitz ML, Papathanasiou S, Doerfler PA, et al. Chromothripsis as an on-target consequence of CRISPR-Cas9 genome editing. Nat Genet 2021;53:895–905.

