## Supplemental Figure S1 for "Optimization of NLS Composition Improves CRISPR-Cas12a Editing Rates in Human Primary Cells"

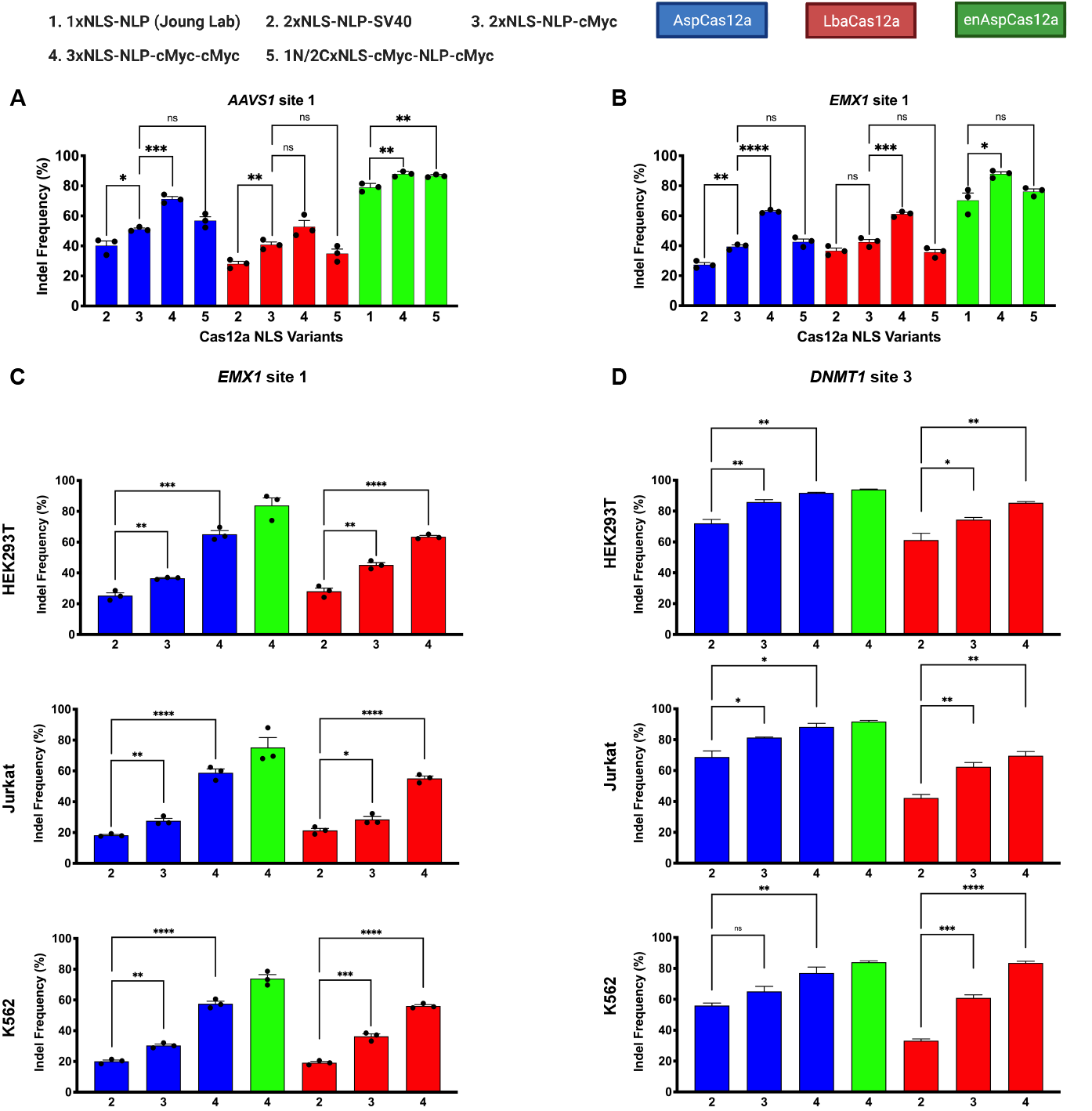

***Supplementary Figure S1. Characterization of Cas12a NLS variant editing in mammalian cell lines.*** (A) Quantification of editing efficiency by Cas12a NLS variant RNPs targeting *AAVS1* site 1 in HEK293T cells at 5 pmol of protein:crRNA complex when delivered by nucleofection (Neon [Invitrogen]). (B) Quantification of editing efficiency by Cas12a NLS variant RNPs targeting *EMX1* site 1 in HEK293T cells at 5 pmol of protein:crRNA complex when delivered by nucleofection. (C) Quantification of editing efficiency by Cas12a NLS variant RNPs targeting *EMX1* site 1 in HEK293T, Jurkat, and K562 cells at 5 pmol of protein:crRNA complex when delivered by nucleofection. (D) Quantification of editing efficiency by Cas12a NLS variant RNPs targeting *DNMT1* site 3 in HEK293T, Jurkat, and K562 cells at 5 pmol of protein:crRNA complex when delivered by nucleofection. Results were obtained from three independent biological replicates and presented as mean ±s.e.m. Statistical significance is determined by two-tailed Student's *t*-test: *ns,* P > 0.05; *, P < 0.01; **, P < 0.001; ***, P < 0.0001; ****, P < 0.00001.
