## Supplemental Figure S2 for "Optimization of NLS Composition Improves CRISPR-Cas12a Editing Rates in Human Primary Cells"

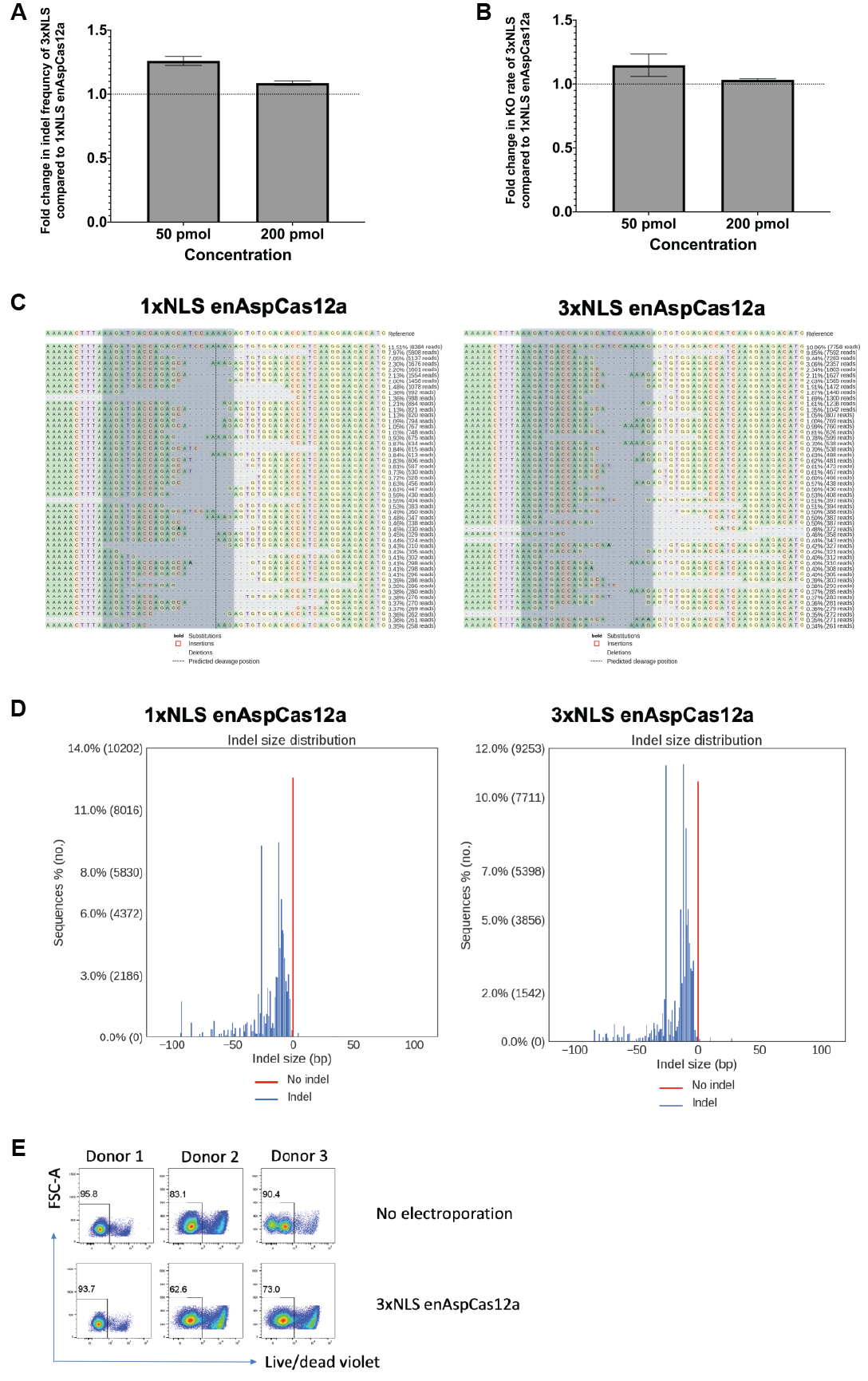

***Supplementary Figure S2. Robust genome editing by enAspCas12a NLS variants in NK cells at IFNG.*** (A) Fold change analysis for editing activity comparing 1xNLS to 3xNLS enAspCas12a RNPs targeting the *IFNG* target site. (B) Fold change analysis in Knockout rate comparing 1xNLS to 3xNLS enAspCas12a RNPs targeting the *IFNG* target site. NK cells were electroporated with 2 doses (50pmol or 200pmol RNP). %Knockout in NK cells stimulated for 3 hours (PMA+ionomycin) or overnight (IL-12+IL-15+IL-18 cocktail). % Knockout = (% IFNG+ cells of non-target group - % IFNG+ cells in experimental group)/% IFNG+ cells of non-target group. (C) Representative sample of the allele frequency table around the protospacer for 1xNLS *(Left)* and 3xNLS *(Right)* enAspCas12a RNP targeting the *IFNG* target site. Blue box highlights the spacer sequence. (D) Representative Indel distribution length for 1xNLS *(Left)* and 3xNLS *(Right)* enAspCas12a RNPs targeting the *IFNG* target site. (E) A representative flow cytometry plot on the cells for Live/Dead staining. Cells were stained with Live and Dead violet viability kit.
