## Supplemental Figure S3 for "Optimization of NLS Composition Improves CRISPR-Cas12a Editing Rates in Human Primary Cells"

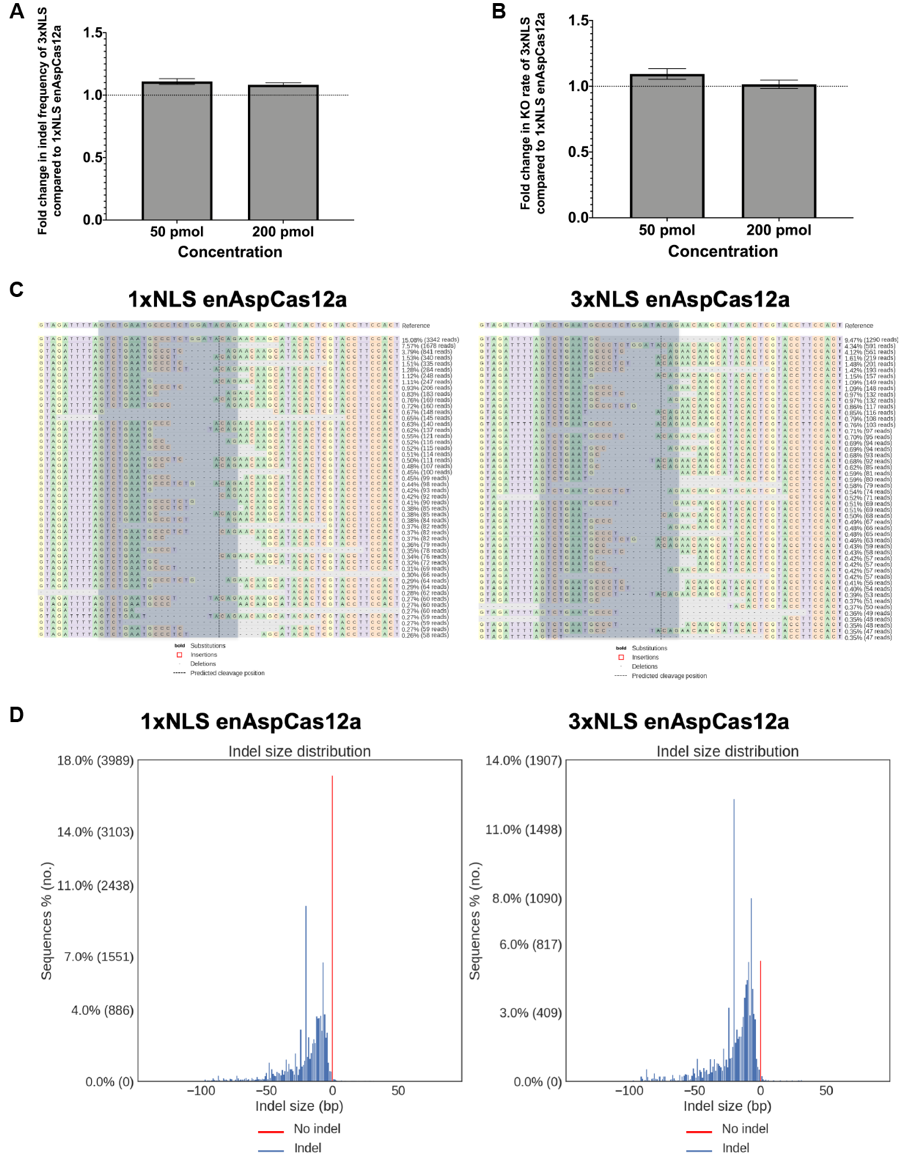

***Supplementary Figure S3. Robust genome editing by enAspCas12a NLS variants in NK cells at CD96.*** (A) Fold change analysis for editing activity comparing 1xNLS to 3xNLS enAspCas12a RNPs targeting the *CD96* target site. (B) Fold change analysis in knockout rate comparing 1xNLS to 3xNLS enAspCas12a RNPs targeting the *CD96* target site. NK cells were electroporated with 2 doses (50pmol or 200pmol RNP). %Knockout in NK cells stimulated for 3 hours (PMA+ionomycin) or overnight (IL-12+IL-15+IL-18 cocktail). % Knockout = (% CD96+ cells of non-target group - % CD96+ cells in experimental group)/% CD96+ cells of non-target group. (C) Representative sample of the alleles frequency table around the protospacer for 1xNLS *(Left)* and 3xNLS *(Right)* enAspCas12a RNPs targeting the *CD96* target site. Blue box highlights the protospacer sequence. (D) Representative Indel distribution length for 1xNLS *(Left)* and 3xNLS *(Right)* enAspCas12a targeting the *CD96* target site.
