## Supplemental Figure S4 for "Optimization of NLS Composition Improves CRISPR-Cas12a Editing Rates in Human Primary Cells"

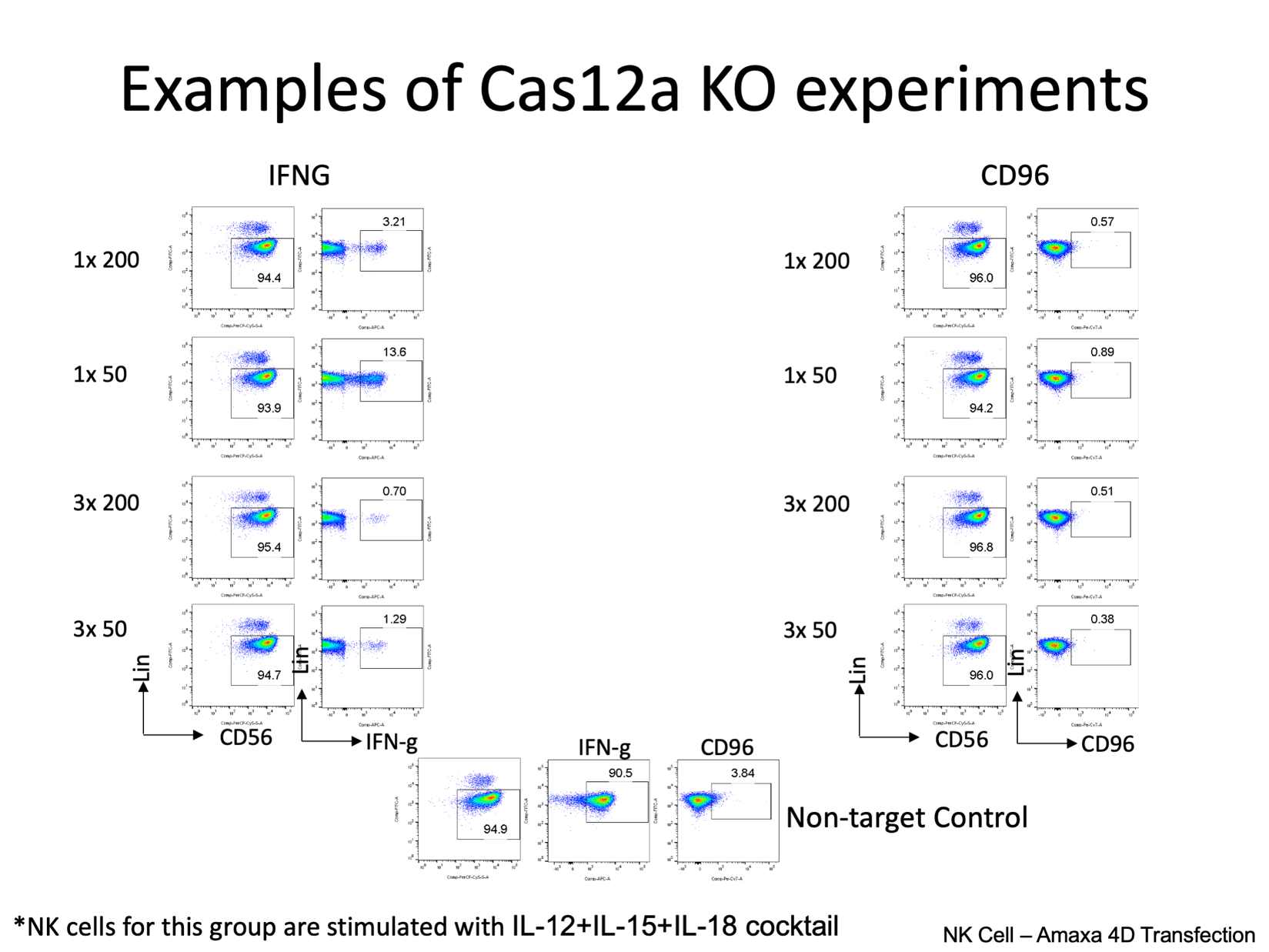

***Supplementary Figure S4. A representative flow cytometry plot on the cells expressing IFN-y or CD96.*** Cells were stained with anti-Lin and anti-CD56 antibodies to gate for NK cells and then stained with anti-IFNy and anti-CD96 antibodies to detect expression of the target protein. Box represent populations of interest. Non-target control is an RNP targeting DNMT1S3.
