## Supplemental Figure S5 for "Optimization of NLS Composition Improves CRISPR-Cas12a Editing Rates in Human Primary Cells"

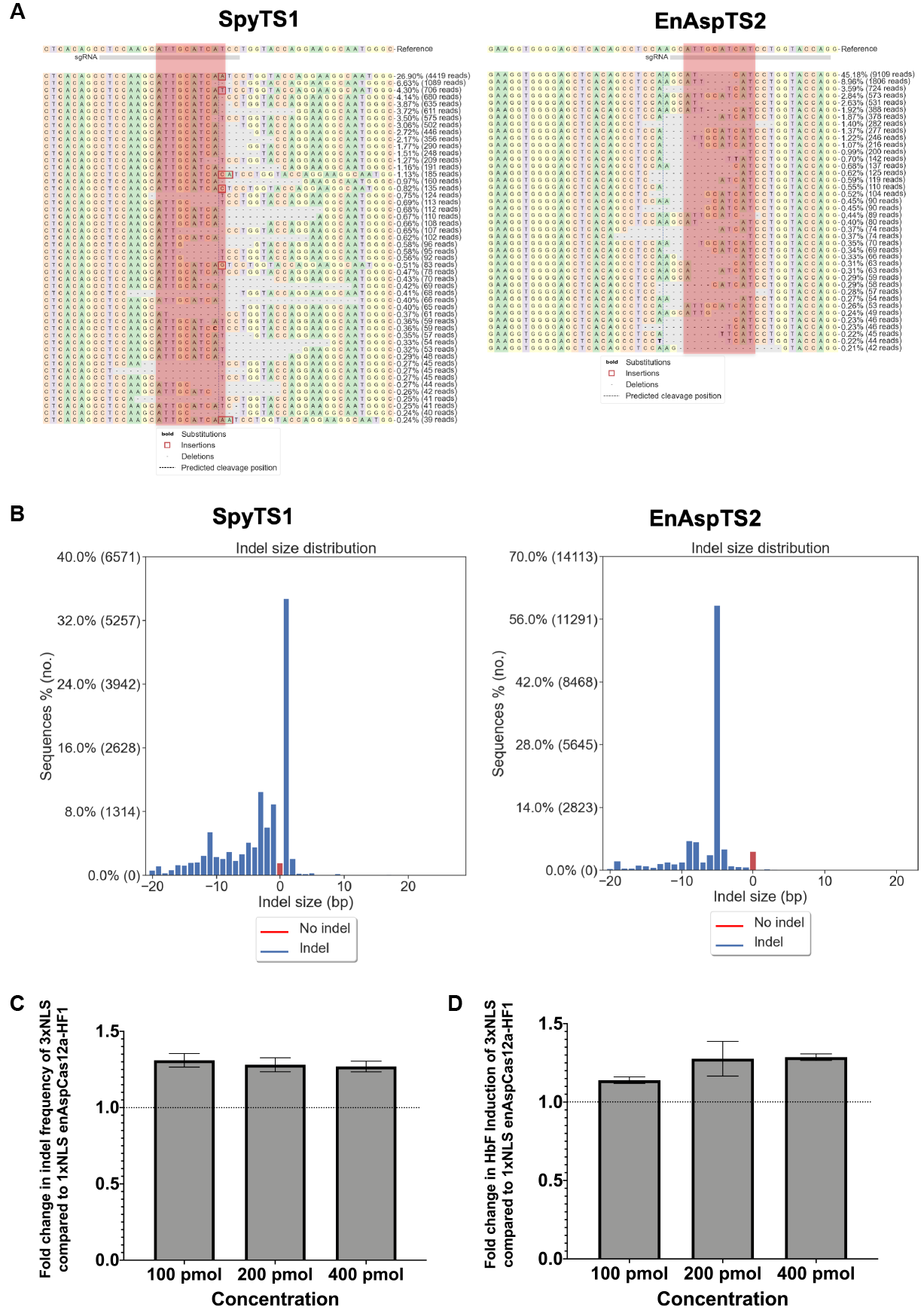

***Supplementary Figure S5. Robust genome editing by SpyCas9 and enAspCas12a at ATF4-binding motif within the BCL11A erythroid-lineage-specific +55 kb enhancer.*** (A) Representative samples of the alleles frequency table around the protospacer for SpyTS1 *(Left)* and EnAspTS2 *(Right)*. Red box highlights the ATF4 binding-motif. (B) Representative Indel distribution for SpyTS1 *(Left)* and EnAspTS2 *(Right)* editing. (C) Fold change analysis for editing activity in CD34+ HSPCs comparing 1xNLS to 3xNLS enAspCas12a-HF1 RNPs targeting EnAspTS2 in titration experiment. (D) Fold change analysis for HbF induction comparing 1xNLS to 3xNLS enAspCas12a-HF1 RNPs targeting EnAspTS2.
