## Supplemental Figure S6 for "Optimization of NLS Composition Improves CRISPR-Cas12a Editing Rates in Human Primary Cells"

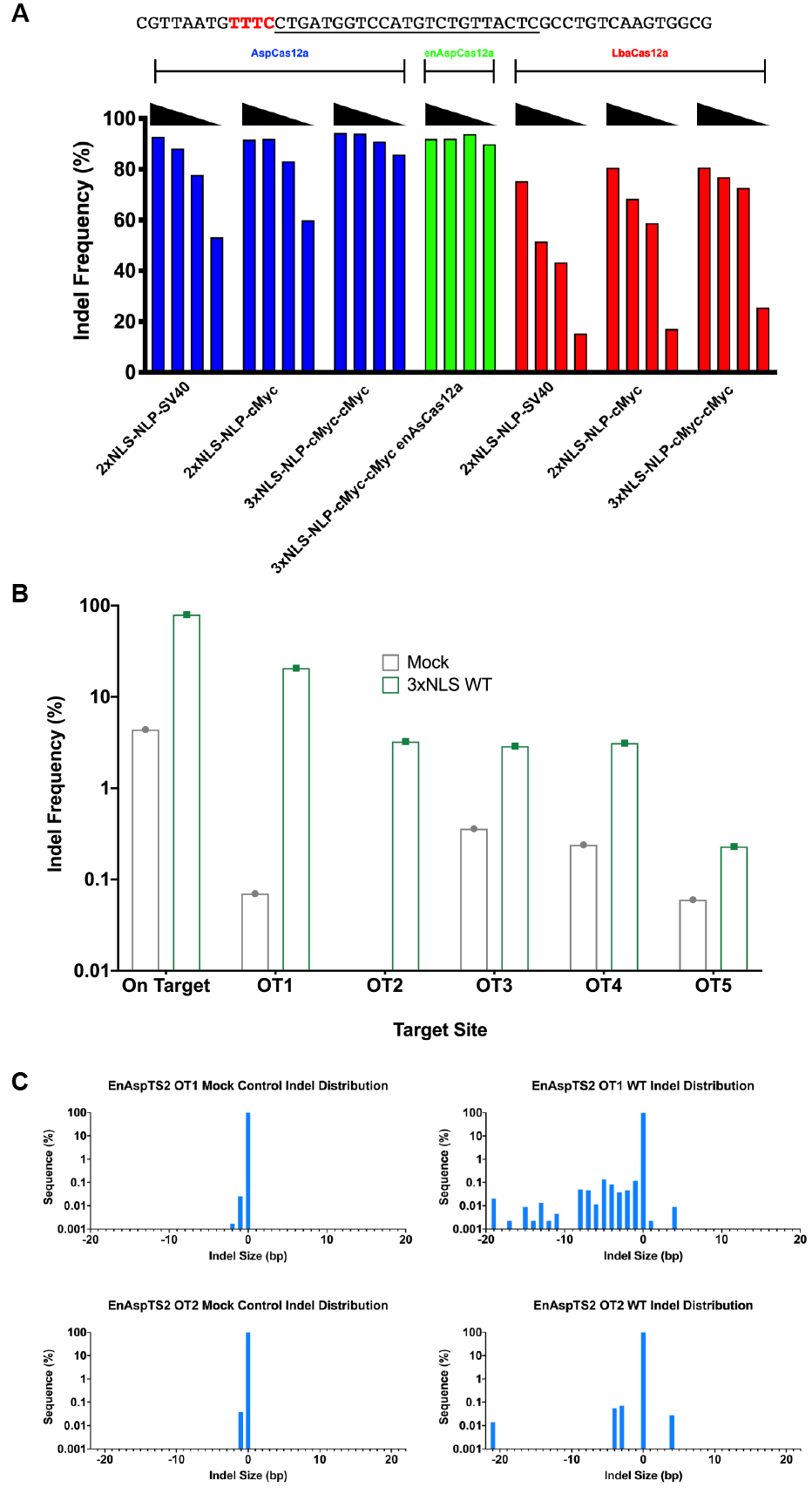

***Supplementary Figure S6. Specificity of Cas12a NLS variants in HEK293T cells at DNMT1 site 3 and CD34+ HSPCs at EnAspTS2.*** (A) Quantification of the nuclease activity of different NLS variants at different Cas12a RNP concentrations (20, 10, 5, and 2.5 pmol of protein:crRNA complex) at *DNMT1* site 3 when delivered by nucleofection to HEK293T cells to determine optimal concentration to produce similar editing rates at the on-target site for specificity analysis. Black triangle indicates the titration of the RNP complex. (B) Quantification of editing activity by enAspCas12a variants at on-target and top 5 GUIDE-tag identified potential off-target sites in HEK293T cells. Results were obtained from a single experiment. (C) Representative Indel distributions by mock control *(Left Column)* and enAspCas12a-WT RNP *(Right Column)* at OT1 *(Top Row)* and OT2 *(Bottom Row)* of EnAspTS2.
